# Chromosome-level and haplotype-resolved genome assembly enabled by high-throughput single-cell sequencing of gamete genomes

**DOI:** 10.1101/2020.04.24.060046

**Authors:** José A. Campoy, Hequan Sun, Manish Goel, Wen-Biao Jiao, Kat Folz-Donahue, Nan Wang, Manuel Rubio, Chang Liu, Christian Kukat, David Ruiz, Bruno Huettel, Korbinian Schneeberger

**Affiliations:** Department of Chromosome Biology, Max Planck Institute for Plant Breeding Research, Carl-von-Linné-Weg 10, 50829 Cologne, Germany; Faculty of Biology, LMU Munich, Großhaderner Str. 2, 82152 Planegg-Martinsried, Germany; FACS & Imaging Core Facility, Max Planck Institute for Biology of Ageing, 50931 Cologne, Germany; Center for Plant Molecular Biology (ZMBP), University of Tübingen, Auf der Morgenstelle 32, 72076 Tübingen, Germany; Departament of Plant Breeding, CEBAS-CSIC, PO Box 164, E-30100, Espinardo, Murcia, Spain; Institute of Biology, University of Hohenheim, Garbenstraße 30, 70599 Stuttgart, Germany; Max Planck-Genome-center Cologne, Carl-von-Linné-Weg 10, 50829 Cologne, Germany

**Keywords:** single-cell sequencing, haplotype-resolved assembly, haplotyping, phasing, *de novo* assembly

## Abstract

Generating chromosome-level, haplotype-resolved assemblies of heterozygous genomes remains challenging. To address this, we developed gamete binning, a method based on single-cell sequencing of haploid gametes enabling separation of the whole-genome sequencing reads into haplotype-specific reads sets. After assembling the reads of each haplotype, the contigs are scaffolded to chromosome-level using a genetic map derived from the gametes. As a proof-of-concept, we assembled the two genomes of a diploid apricot tree based on whole-genome sequencing of 445 individual pollen grains. The two haplotype assemblies (N50: 25.5 and 25.8 Mb) featured a haplotyping precision of >99% and were accurately scaffolded to chromosome-level.

## Introduction

Currently, most diploid genome assemblies ignore the differences between the homologous chromosomes and assemble the genomes into one pseudo-haploid sequence, which is an artificial consensus of both haplotypes. Such an artificial consensus can result in imprecise gene annotation and misleading biological interpretation^1,2^. To avoid these problems, it is a common strategy to inbreed or to generate double-haploid genotypes to enable the assembly of homozygous genomes.

Recent alternatives allowing for the assembly of both haplotypes include chromosome sorting^3^, Strand-seq^4–6^ and high-throughput chromosome conformation capture (Hi-C)^7–13^ sequencing. Chromosome sorting separates individual chromosomes before sequencing, and thus enables the sequencing and assembly of individual haplotypes. However, sorting of particular chromosomes may not always be possible if they cannot be discriminated based on their fluorescence intensity or light scatter^14^ and may need tedious generation of specific lines for sorting^15^. The more recent method Strand-seq is a single-cell technique that requires neither parents nor gametes which can be potentially used to cluster long sequencing reads by chromosome, phase haplotypes, and scaffold using genetic map techniques, however, the difficulty for generating Strand-seq data has limited its application to a narrow number of model species. In contrast, the analysis of the chromosome conformation, including Hi-C technologies which enable the detection of chromatin interactions at an unprecedented scale, has been successfully applied for haplotype phasing and genome scaffolding for a wide range of species ^7,9–12,16^. However, despite its simple application, Hi-C-based phasing can be error prone due to some weaknesses in defining the alleles that distinguish haplotypes, which in turn can lead to haplotype switch errors^11^ and result in mis-scaffolding of small contigs due to the lack of sufficient informative connections to other contigs^8,12,13^. Also the reconstruction of whole chromosomes structures can be error-prone as already one local mis-scaffolding is sufficient to introduce severe mis-assemblies like falsely joining chromosome arms^10^. It is therefore necessary to carefully inspect assemblies that rely on Hi-C for phasing or scaffolding to identify errors, which in turn require correction based on additional evidence including, for example, the integration of genetic maps ^8,10,17^.

An elegant alternative for haplotype phasing, called trio binning, is based on the separation of whole-genome sequencing reads into haplotype-specific read sets before assembly using the genomic differences between the parental genomes^2^. While this is a powerful method, it can be limiting if the parents are not available or are unknown^18^. A solution for this is the sequencing of a few gamete genomes (derived from the focal individual), which is sufficient for the inference of genome-wide haplotypes, but relies on existing long-contiguity reference sequences^19–22^.

In addition to resolving haplotypes, the generation of chromosome-level assemblies, which are necessary to understand the full complexity of genomic differences including all kinds of structural rearrangements, is similarly challenging^23,24^. While recent improvements in long DNA molecule sequencing^25^ and as mentioned above in Hi-C data generation promise the assembly of telomere-to-telomere contigs, genetic maps can reliably help to resolve mis-assemblies and guide chromosome-level scaffolding^10^. The generation of genetic maps, however, relies on a substantial amount of meiotic recombination which usually implies the genotyping of hundreds of recombinant genomes^26,27^. Creating and genotyping sufficiently large populations is not possible in some species (like many of the mammals including humans), and for those species for which it is possible it can be time-consuming and costly, and may post great challenges if the individuals show long juvenility or sterility^16^.

To address all these challenges, we present gamete binning, a method for chromosome-level, haplotype-resolved genome assembly - independent of parental genomes or recombinant progenies (Fig. 1). The method starts by isolating gamete nuclei from the focal individual followed by high-throughput single-cell sequencing of hundreds of the haploid gamete genomes. (For clarification, we collectively refer to both gametophytes in plants and gametes in animals collectively as gametes, as both have haploid genomes.) The segregation of sequence variation in the gamete genomes enables a straightforward phasing of all variants into two haplotypes, which subsequently allows for genetic mapping and sorting of whole-genome sequencing reads into distinct read sets - each representing a different haplotype. Assembling these independent read sets leads to haplotype-resolved genome assemblies, which can be scaffolded to chromosome-level using a gamete genome-derived genetic map.

**Figure 1.**
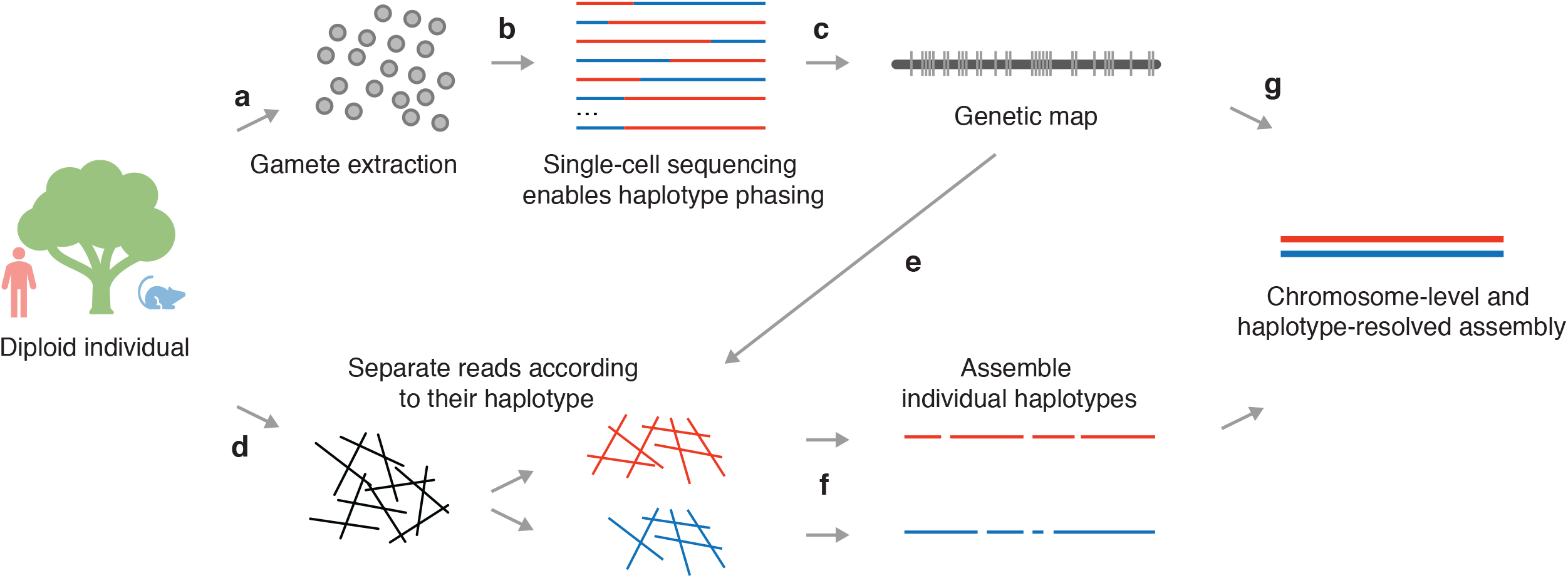
Overview of gamete binning. **a**. Extraction of gamete nuclei. **b**. Single-cell genome sequencing of haploid gametes and haplotype phasing. **c**. Genetic map construction based on the recombination patterns in the gamete genomes. **d**. Long-read sequencing of somatic material. **e**. Separation of long reads based on genetic linkage groups using phased alleles. **f**. Independent assembly of each haplotype of each linkage group. **g**. Scaffolding assemblies to chromosome-level using the gamete-derived genetic map.

## Results

### Preliminary diploid-genome assembly

We used gamete binning to assemble the two haploid genomes of a specific, diploid apricot tree (*Prunus armeniaca*; cultivar ‘Rojo Pasion’^28^), which grows in Murcia, southeastern Spain (Supplementary Figure 1). We first performed a preliminary *de novo* genome assembly using *Canu*^29^ with 19.9 Gb long reads (PacBio, Supplementary Figure 2) derived from DNA extracted from fruits and corresponding to 82x genome coverage according to a genome size of ∼242.5 Mb estimated by *findGSE*^30^ (Methods; Supplementary Figure 3). After purging haplotype-specific contigs, the curated assembly consisted of 939 contigs with a combined length of 230.9 Mb and an N50 of 563.8 kb, which represents a haploid, but mosaic assembly of the apricot genome (Methods).

**Figure 2.**
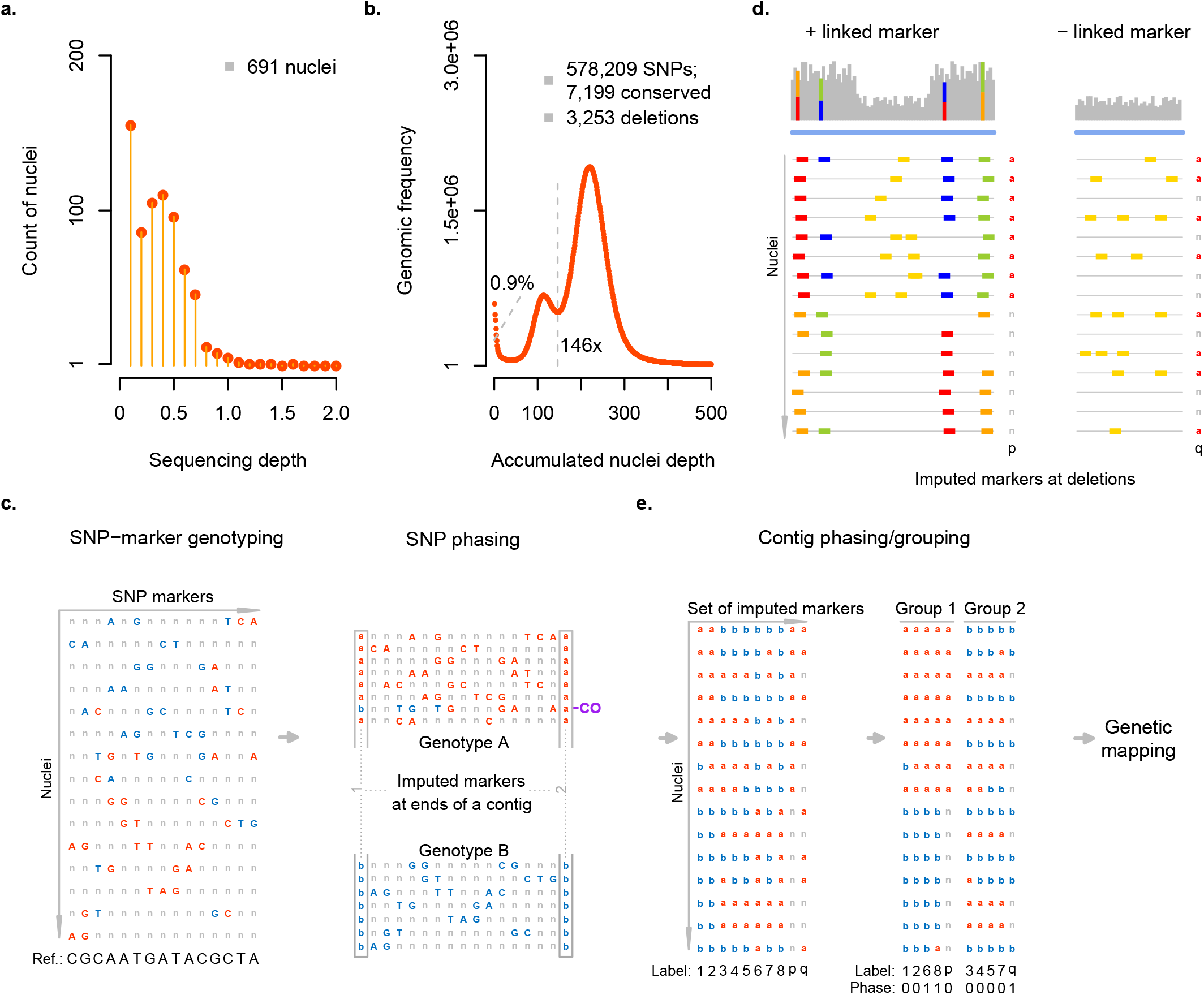
Single-pollen nuclei sequencing, variant phasing and genetic mapping. **a**. Sequencing depths of 691 pollen nuclei. **b**. Sequencing depth histogram of pooled pollen short reads. The left-most peak revealed 0.9% of the genome that were not well covered in the pollen read sets (i.e., ≤5x). The middle peak indicated regions covered only by half of the genomes and present in only one of the haplotypes, and the right-most peak indicated regions, which were present in both haplotypes and showed the expected coverage. In regions represented in both haplotypes, 578,209 SNPs were defined. Regions without SNP markers were classified into 3,253 deletions and 7,199 conserved regions (Methods). **c**. SNP phasing along contigs. Genotyping was first performed for each individual nuclei at each SNP marker. As shown, both genotypes (in red and blue) were mixed in the curated but mosaic assembly. After phasing, 8 and 7 nuclei were respectively clustered for genotype A and B, and crossover could be identified. With this, representative markers were imputed at ends of contigs. **d**. Imputation of markers at deletions by genotyping using normalized read count. Two cases were considered for phasing (and positioning) a deletion marker (in the genetic map). If it was linked with surrounding SNP alleles, it could be phased accordingly; otherwise, comparison its genotype sequence to genotype sequences of all other markers (including SNP-derived markers at ends of contigs) would be performed to find its value of phase (and positioning). **e**. Linkage group and genetic map construction using the set of imputed markers (SNP-derived markers labeled as 1-8 and deletion markers as *p* and *q*). For example, the genotype sequences of 6, 8 and *q* needed to be flipped (i.e., phase values were 1 - contig phasing). Further ordering of the markers (using *JoinMap*) led to linkage group-wise genetic maps.

**Figure 3.**
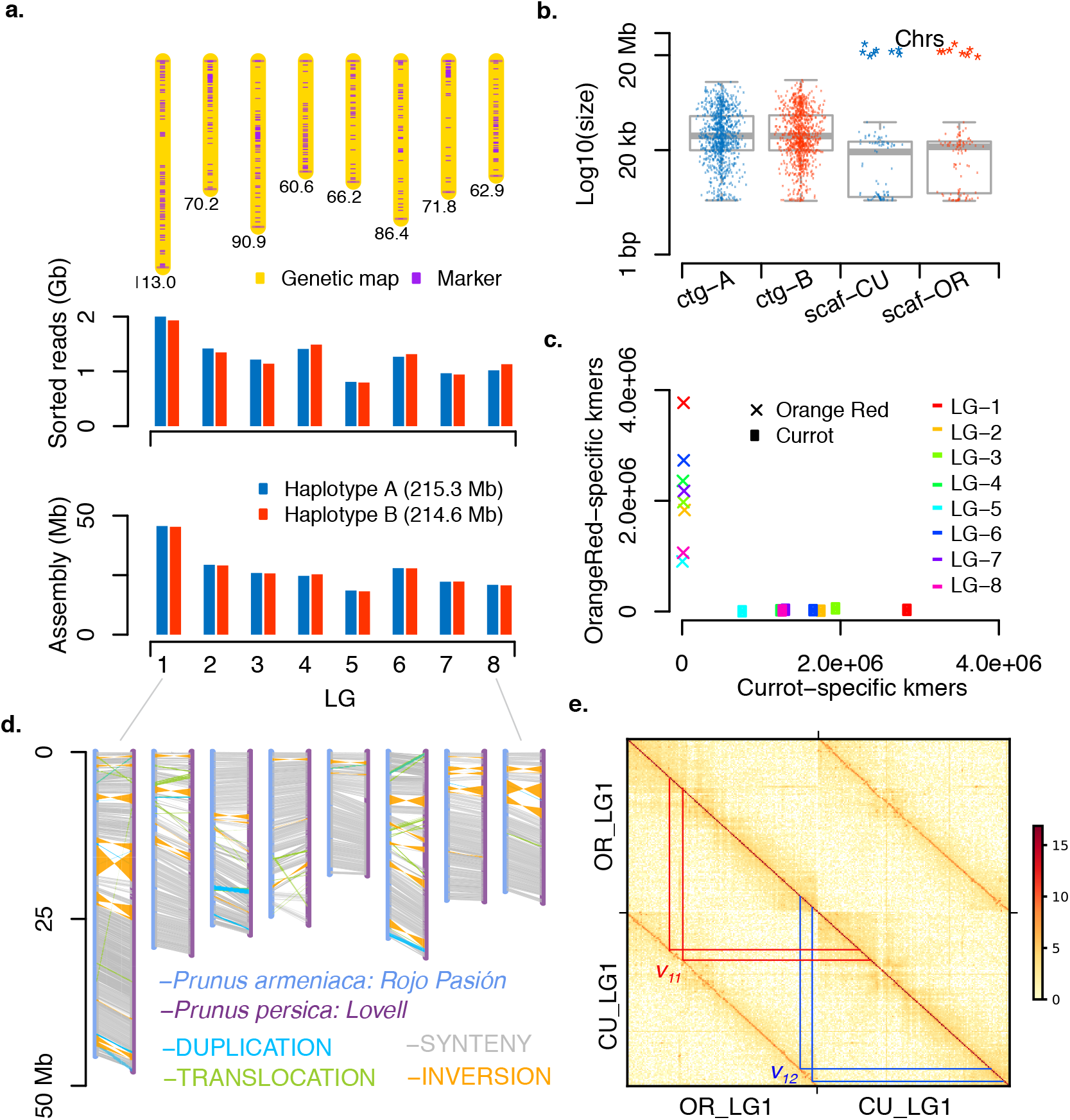
Genetic mapping, haplotype-specific assembly and validation. **a**. Top: Genetic map with a total genetic length of 622.0 cM (Methods). Middle: up to 2 Gb reads were assigned to one of the two haplotypes of each linkage group. Bottom: a combination of haplotype-A/B linkage groups led to two assemblies with 214.6 and 215.3 Mb. **b**. Contig size distributions before (ctg-A, ctg-B) and after scaffolding (scaf-CU for the assembly with sequence from ‘Currot’; and scaf-OR for the assembly with sequence from ‘Orange Red’). After scaffolding, eight chromosome-scale pseudo-molecules were obtained for each haplotype as labeled by “Chrs”. **c**. Haplotype validation for the two assemblies of each linkage group (LG-1-8) using parent-specific *k*-mers (of ‘Orange Red’ and ‘Currot’). With each linkage group, the two assemblies could be clearly identified as either ‘Currot’-haplotype or ‘Orange Red’-haplotype using parental *k*-mers. After combining the ‘Currot’-related assemblies and ‘Orange Red’-related assemblies to genome-level, *k*-mer comparison revealed a haplotype accuracy of 99.1%. **d**. Using the ‘Currot’-haplotype as representative and comparing it to the assembly of the double haploid Prunus ssp. reference genome (*Prunus persica*, and other closely-related species; Supplementary Figure 6) revealed high levels of synteny and thus implies high accuracy of the genetic map and chromosome-level scaffolding. **e**. Hi-C contact map based on the assemblies of the two haplotypes of chromosome 1 (Currot (CU) and Orange Red (OR)) at a resolution of 300 kb. The contact signal showed a high contiguity within the haplotypes (main diagonal line) and confirmed two large inversions (v_11_ and v_12_) which we observed in the assembled sequence of the two haplotypes. (See Supplementary figures 9-15 for chromosomes 2-8).

### High-throughput single-cell sequencing of pollen

To advance this assembly, we isolated pollen grains from ten closed flowers (to avoid contamination of foreign pollen) and released their nuclei following a protocol based on pre-filtering followed by bursting^31^ (Fig. 1a; Methods). The nuclei mixture was cleaned up using propidium iodide staining plus sorting by flow cytometry, leading to a solution with 12,600 nuclei that were loaded into a 10x Chromium Controller in two batches - each with 6,300 nuclei (Supplementary Figures 1a-d; Supplementary Figure 4; Methods). With this we generated two 10x single-cell genome (CNV) sequencing libraries, which were sequenced with 95 and 124 million 151 bp paired-end reads (Illumina). By exploring the *cellranger*-corrected cell barcodes within the read data of both libraries, we extracted 691 read sets - each with a minimum of 5,000 read pairs (Methods; Fig. 2a).

**Figure 4.**
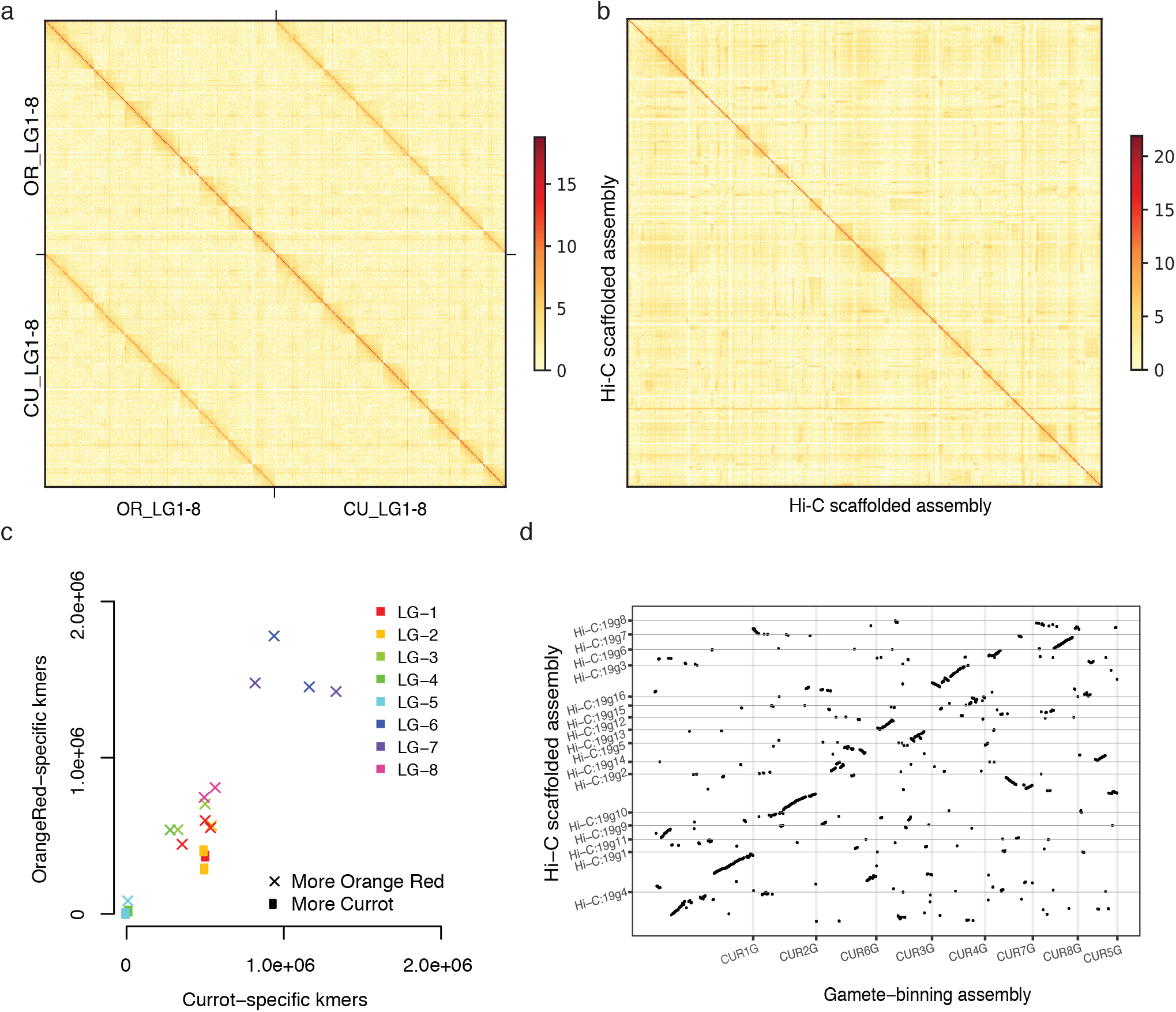
Comparison of Hi-C based phasing and scaffolding with gamete binning. **a**. Hi-C contact map based on all 16 haplotype assemblies generated with gamete binning (Currot (CU) and Orange Red (OR)). **b**. Hi-C contact map based on all 16 haplotype assemblies generated with Hi-C data (Currot (CU) and Orange Red (OR)). Note the contact signal along the main diagonal was much weaker as compared to the signal based on the gamete binning assembly, and virtually no contact signals between two different haplotypes could be identified. **c**. Haplotype validation of each linkage group (LG-1-8) of the Hi-C assembly using parent-specific *k*-mers. Almost all haplotype-specific assemblies included *k-*mers specific to both of the parental alleles indicating severe errors in the phasing. **d**. Alignment of the Hi-C based assembly to genetic map derived assembly (i.e. gamete binning derived assembly) revealed mis-joining or splitting of linkage groups within the Hi-C assembly. For example, CUR1G was split into Hi-C:19g1, 19g4; on the other hand, alignments of 19g2 to CUR2G, CUR5G and CUR7G revealed mis-joins of sequences from independent linkage groups.

Aligning the reads of each pollen genome to the curated assembly, we found that the reads of 246 sets featured high similarity to thrip genomes or included more than one haploid genome, possibly due to random attachment of multiple nuclei during 10x Genomics library preparation or the uncompleted separation of pollen nuclei during pollen maturation^32^ (Supplementary Figure 6a-c; Methods). Thus, we selected a set of 445 haploid pollen genomes. In general, the short-read alignments did not show any biases or preferences for specific regions of the genome as reported for some of the single-cell genome amplification kits, but covered nearly all regions (99.1%) of the curated assembly (Fig. 2b; Supplementary Figure 6d).

### Haplotype phasing and genetic mapping

With short read alignments, we identified 578,209 heterozygous SNPs on 702 contigs with a total length of 218.0 Mb (Fig. 2b; Methods). Even though this implied 1 SNP marker every 377 bp on average, we observed that the distances between some of the SNP markers were larger than the usual long reads, which would hamper the haplotype assignment of reads whenever they aligned to such regions. Overall, we observed 10,452 regions larger than 2 kb without markers (110.9 Mb) including 237 regions (12.5 Mb), that spanned entire contigs. Regions without markers occur if the two haplotypes are identical (which is a common phenomenon in domesticated genomes) or if a region exists only in one of the haplotypes (e.g. a large indel). We distinguished these two cases using the short-read coverage of the combined pollen read sets, assuming that the regions that are only present in one haplotype are supported by only approximately half of the reads (Methods). While 7,199 regions (74.5 Mb) were shared between the haplotypes (and were labelled as conserved), we found that 3,253 regions (36.4 Mb) were specific to one of the haplotypes (i.e. deletions; Fig. 2b). Such regions (i.e. deletions) which are specific to one haplotype can also be used as markers. If such deletions were linked to nearby SNP markers, we phased them according to their linked alleles. For deletions on contigs without additional markers, we used the absence and presence of read alignments in the pollen to assign genotypes.

The haploid nature of the 445 selected individual pollen genomes allowed us to phase all SNP and deletion markers into two haplotypes simply by using the linkage within the pollen genomes (Fig. 2c-d). To phase the haplotypes across the contigs, we generated two virtual markers for each contig representing the (imputed) alleles at both ends of the contig. The markers were grouped into a genetic map with eight linkage groups (corresponding to the eight homologous chromosome pairs) including 891 contigs with a total length of 228.0 Mb (corresponding to about 99% of the complete assembly) using *JoinMap* 4.0^33^ (Fig. 2e; Fig. 3a) (Methods).

### Haplotype-specific long read separation and chromosome-level assembly

After this, we aligned the PacBio reads to the curated assembly. Using the phased alleles (of the SNP and deletion markers) within each of the individual PacBio read alignments, we separated 93.4% of the reads into one of 16 haplotype-specific clusters representing the two haplotypes of each of the eight linkage groups. Reads that aligned in regions that were conserved between the two haplotypes were randomly assigned to one of the two haplotype-specific clusters (Fig. 3a; Methods). Similarity analyses revealed that most of the remaining 6.6% reads were related to organellar genomes or repetitive sequences.

The 16 haplotype-specific read sets were independently assembled using *Flye*^34^, which led to 16 haplotype-specific chromosome assemblies with average N50 values ranging from 662.3 to 664.6 kb (Table 1; Methods). Using the genetic map, we combined the contigs of each assembly into a pseudo-molecule. This led to two haplotype-resolved chromosome-level assemblies, both with N50 above 25.0 Mb (Fig. 3a-b; Methods).

**Table 1.** Assembly and validation statistics of two haplotype-resolved genome assemblies. Note, the eight main chromosome-level scaffolds of each haplotype made up ∼99% of the respective assembly.

| Haploid assemblies<br>of 'Rojo Pasion' | Genome-specific *k*-mers<br>common with parental WGS |  | Precision in<br>haplotyping | Size<br>[Mb] | Chromosome<br>scaffolds | Contig N50<br>[Mb] | N50 [Mb] | Protein-coding<br>genes<br>(Total genes) |
| --- | --- | --- | --- | --- | --- | --- | --- | --- |
|  | 'Currot' | 'Orange Red' |  |  |  |  |  |  |
| 'Currot'-haplotype | 12,983,934 | 129,874 | 99.1% | 216.0 | 8 | 0.662 | 25.8 | 30,661<br>(52,472) |
| 'Orange Red'-<br>haplotype | 81,422 | 16,807,958 | 99.5% | 215.2 | 8 | 0.664 | 25.5 | 30,378<br>(51,701) |

To assess haplotype accuracy, we additionally whole-genome sequenced the parental cultivars of ‘Rojo Pasion’ known as ‘Currot’ and ‘Orange Red’. Using Illumina sequencing technology, we generated 15.7 and 16.2 Gb short reads of each of the diploid parental genomes, respectively. Overall, we found that ∼99.1% of the *k*-mers that were specific to one of the haplotype assemblies could be found in the corresponding parental genome illustrating that almost all of the variation was correctly assigned to haplotypes (Fig. 3c; Table 1; Methods). Having proved the haplotype accuracy, the assemblies were polished resulting in final haplotype assemblies. The final haplotype assembly sizes were 216.0 and 215.2 Mb for ‘Currot’-genotype (8 scaffolds, N50: 25.8 Mb) and ‘Orange Red’-genotype (8 scaffolds, N50: 25.5 Mb), respectively (Table 1).

We estimated the overall assembly quality by comparing the *k*-mer distributions of the assemblies and the Illumina short read sets of the focal and parental using KAT^35^ and Merqury^36^. Both haplotype genome assembly showed very high quality values (*QV* > 36) and the absence of allelic duplications between the haplotypes, though a fraction of ∼7% of the heterozygous *k*-mers in the reads was missing in the assemblies (Supplementary Figures 7c, 8).

To further assess the overall structures of the assembled chromosomes, we compared them to recently assembled chromosomes of very closely-related species such as the heterozygous ‘Chuanzhihong’ apricot (*Prunus armeniaca*)^37^, the Japanese apricot (*Prunus mume*)^38^, and a more distantly-related species, peach (*Prunus persica:* doubled-haploid genome)^39^ using *SyRI*^23^ (a tool designed for the comparison of chromosome-level assemblies). Our assemblies showed high consistency in the synteny to these assemblies across entire chromosomes, reflecting the reliability of the genetic map and the assembled genome structures (Fig. 3d; Supplementary Figure 6).

As yet another way to assess the quality of the genome we generated two Hi-C libraries from DNA extracted from leaves of Rojo Pasion and sequenced them totaling in 191.2 million read pairs (or ∼240x genome coverage). We created Hi-C contact maps using each of the homologous chromosome pairs separately as well as using the entire genome (Fig. 3e; Fig. 4a; Supplementary Figures 9-15). In general, the contiguity of contact signals surrounding the main diagonal of the map again demonstrated the high quality of the structure of the assemblies.

### Comparing gamete binning with Hi-C based phasing and genome scaffolding

However, the perhaps more interesting way to use the Hi-C data is its application for genome phasing and scaffolding and the comparison of its assembly performance to that of gamete binning.

Applying *ALLHiC*^8^ to the Hi-C reads sets generated 16 scaffolds (representing the 16 haploid chromosomes), with sizes ranging from 11.2 to 51.1 Mb (Methods). (Using a different Hi-C-based phasing and scaffolding tool, *SALSA2*^40^, did not lead to comparable results, thus not compared further.). For comparison, we also generated Hi-C contact maps for *ALLHiC* based assemblies (Fig. 4b). Interestingly, the contact maps of the gamete binning and *ALLHiC* based assemblies were strikingly different. Only the gamete binning assembly showed (beside the contact within the haplotypes) the expected contact signals between two different haplotypes, which also were reported for other species ^8,41^. The absence of these signals in the Hi-C based assembly suggests that the assembly was falsely merging sequences from different haplotypes and the contigs were likely to be scaffolded in the wrong order.

To test if the Hi-C based assemblies were truly a mixture of the two haplotypes, we checked the presence of parental-specific *k*-mers within each of the 16 haplotype-specific chromosome-level assemblies (Fig. 4c). This revealed that the majority of the haplotype specific assemblies were in fact mixtures of the two haplotypes, which is in great contrast with the high haplotyping accuracy of gamete binning. Finally, a whole-genome alignment of the Hi-C based assembly to the genetic map based assembly of gamete binning revealed many ambiguities between the genetic maps and the Hi-C based assembly within essentially all haplotype-specific chromosome assemblies (Fig. 4d).

Taken together, besides its broad application, Hi-C-based phasing and scaffolding was far from being error-free. Some of the errors combined large pieces from different haplotypes, which resulted in falsely arranged chromosomes and severe phasing errors. Though, gamete-binning might be more tedious in its experimental requirements, the improved assembly quality might justify the additional effort.

### Haplotype diversity and (non-allelic) meiotic recombination

In contrast to conventional diploid genome assemblies where the two haplotypes are merged into one artificial consensus sequence, separate haploid assemblies allow for the analysis of haplotype diversity. Comparing the two haplotype assemblies of ‘Rojo Pasion’ using *SyRI*^23^ allowed us to gain first insights into the haplotype diversity within an individual apricot tree. Despite high levels of synteny, the two assemblies revealed large-scale rearrangements (23 inversions, 1,132 translocation/transpositions and 2,477 distal duplications) between the haplotypes making up more than 15% of the assembled sequence (38.3 and 46.2 Mb in each of assemblies; Supplementary Table 1). Using the Hi-C contact maps (Fig. 3e; Supplementary Figures 9-15), we validated the 17 largest rearrangements (> 500 kb) between the haplotype assemblies. Using a comprehensive RNA-seq dataset sequenced from multiple tissues of ‘Rojo Pasion’ including reproductive buds, vegetative buds, flowers, leaves, fruits (seeds removed) and barks as well as a published apricot RNA-seq dataset^37^, we predicted 30,378 and 30,661 protein-coding genes within each of the haplotypes (with an annotation completeness of 96.4% according to a BUSCO^42^ analysis). Mirroring the huge differences in the sequences, we found the vast amount of 942 and 865 expressed, haplotype-specific genes in each of the haplotypes (Methods; Supplementary Tables 2-3). Such deep insights into the differences between the haplotypes, which are only enabled by chromosome-level and haplotype-resolved assemblies, will generally be of high value for the analysis of agronomically relevant variation.

Moreover, the chromosome-level assemblies also allow for fine-grained analyses of the haploid pollen genomes, which have already undergone recombination during meiosis. Meiotic recombination is the major mechanism to generate novel variation in offspring genomes. During meiosis new haplotypes are formed by sequence exchanges between two homologous chromosomes. To keep chromosome structures intact during such exchanges, it is essential that recombination only occurs in syntenic regions as otherwise large parts of the chromosome can be lost or duplicated in the newly formed molecules. Re-analyzing the 445 pollen nuclei genomes using one of the chromosome-level assemblies as reference, we detected 2,638 meiotic crossover (CO) events (Methods). To improve the resolution of the predicted CO events (6.1 kb), we selected 2,236 CO events detected in 369 nuclei with a sequencing depth above 0.1x genome coverage (Supplementary Table 4). Along the chromosomes, CO events were broadly and positively correlated with the density of protein-coding genes and were almost completely absent in rearranged regions as expected (Fig. 5; Methods). By investigating the fine-scale pattern of short read alignment of each nuclei, we identified six CO events located in rearranged regions (0.3% of 2,236 CO events found in 1.6% of the pollen genomes), which led to stark chromosomal rearrangements. In each of the six chromosomes we found duplicated read coverage and pseudo-heterozygous variation in the regions that were involved in the chromosome rearrangements as induced by the non-allelic CO (Fig. 6). This evidences the existence of non-allelic recombination in pollen genomes and might open up a more detailed view on the actual meiotic recombination patterns as compared to what could be observed in offspring individuals.

**Figure 5.**
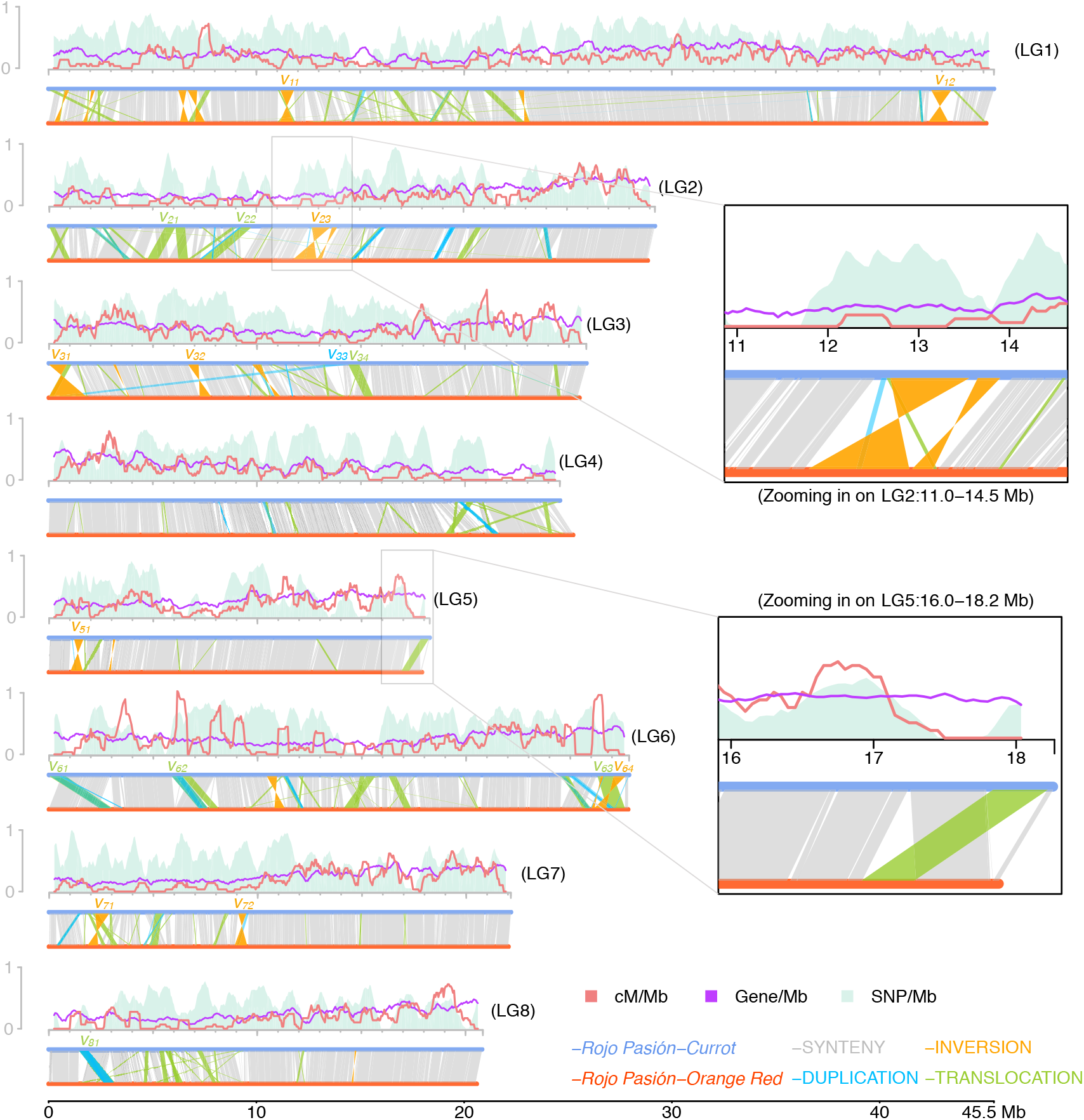
Structural genome variations and meiotic recombination. Top: recombination landscape created with sliding windows of 500 kb at a step of 50 kb with COs detected in all single pollen nuclei (with coverage over 0.1x), coupled with SNP density and gene density. For x-axis, coordinates were based on the haploid assembly of ‘Currot’-genotype. For y-axis, all features were scaled to 1.0, which stands for a maximum of 18 for recombination frequency (*cM/Mb*), 7,410 for SNP density and 480 for gene density. Bottom: structural variations (>50 kb) identified between the two haploid assemblies. In general, crossovers are almost completely absent in SVs, for example, at LG2:11.0−14.5 Mb (inversion case) and LG5:16.0−18.2 Mb (translocation case). Variants spanning over 500 kb are labelled as *v*_*xy*_, where *x* denotes the chromosome number and *y* the identifier of the variant in the chromosome. All these large variants were confirmed within Hi-C contact maps (Fig. 3e, Supplementary Figures 9-16).

**Figure 6.**
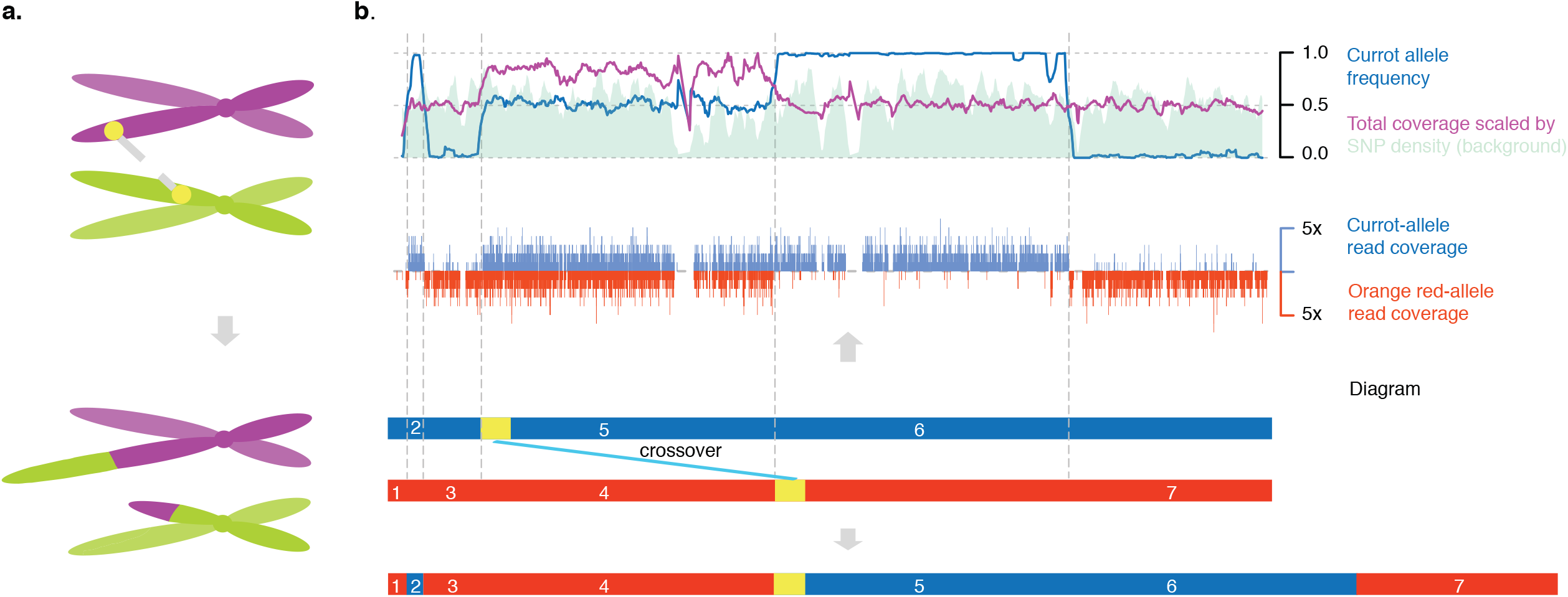
Non-allelic crossovers and its consequences. **a**. Illustration of a non-allelic crossover which results in a chromosomal anomaly. **b**. Analysis of a single-pollen nuclei, which revealed a non-allelic CO resulting in the duplication of a large chromosomal segment. The short-read alignments of a haploid nuclei revealed a pseudo-heterozygous region with increased read coverage, which is the hallmark of a long duplication specific to this genome. All other chromosomes were haploid (not shown). (Top row: ‘Currot’ allele frequency, SNP density (in sliding windows of 500 kb at a step of 50 kb), and read coverage scaled by SNP density. Middle row: count of ‘Currot’ or ‘Orange Red’ alleles at SNP markers. Bottom row: diagram illustrating how a non-allelic CO in transposed regions (as indicated by yellow rectangles) resulted in a large duplication, i.e., the original homologous chromosomal regions labelled with “4” and “5” are now part of the same newly formed chromosome.

## Conclusion

Taken together, following the elegant rationale of haplotype-based read separation before genome assembly introduced by trio binning^2^, we present gamete binning. In contrast to trio binning, gamete binning does not rely on paternal genomes, but instead uses the genomes of individual gametes to resolve haplotypes. In addition, the recombination patterns in these gamete genomes can be used to calculate a genetic map, which in turn enables the generation of chromosome-level assemblies. High-throughput analysis of gamete genomes avoids tedious generation of offspring progeny and allows to sample the required material in its ecological context, which makes it possible to analyze meiotic recombination as it occurs in natural environments. As a result, gamete binning can efficiently and effectively enable haplotype-resolved and chromosome-level genome assembly of any heterozygous individual with accessible gametes.

## Online Methods

### DNA extraction, Illumina/PacBio library preparation and sequencing

Fresh developing fruits of ‘Rojo Pasion’ were frozen in liquid nitrogen immediately after being sampled in Murcia, Spain. After being shipped to the Max Planck Institute for Plant Breeding Research (MPIPZ, Cologne, Germany), DNA was extracted from the mesocarp and exocarp of the fruits using the Plant DNA Kit of Macherey-Nagel™ to create a PacBio sequencing library. Meanwhile, fresh leaves were sampled from the parental cultivars (‘Currot’ and ‘Orange Red’) at the experimental field of CEBAS-CSIC in Murcia, Spain, and Illumina short read libraries were prepared after DNA extraction using the Plant DNA Kit of Macherey-Nagel™.

All libraries were sequenced with the respective sequencing machines (Illumina HiSeq 3000 and PacBio Sequel I) at Max Planck Genome-centre Cologne (MP-GC), which led to 19.9 Gb long reads for ‘Rojo Pasion’ (PacBio; Supplementary Figure 2) and 15.7 and 16.2 Gb short reads for the parental cultivars (Illumina). Note that the parental WGS data were only used for haplotype validation and for sorting the individual chromosome assemblies to two sets of eight chromosomes to match the inheritance of the chromosomes.

### Pollen nuclei DNA extraction, 10x sc-CNV library preparation and sequencing

Dormant shoots of ‘Rojo Pasion’ bearing developed flower buds were collected in Murcia, Spain. Then, the shoots were shipped at 4 °C to MPIPZ (Cologne, Germany) and were grown in long-day conditions in the greenhouse. Flowers at the pre-anthesis stage were frozen in liquid nitrogen. Anthers from ten ‘Rojo Pasion’^28^ flowers were extracted with forceps and submerged in woody pollen buffer (WPB)^43^. Around 500,000 pollen grains were extracted from anthers by vortexing them in WPB. The nuclei were isolated from the pollen using a modified bursting method^31^. Isolated pollen was prefiltered (100μm) and bursted (30um) using Celltrics™ sieves and woody pollen buffer. The nuclei were then stained with propidium iodide (PI) at 50 μg/mL just before sorting and counting by flow cytometry to remove pollen grain debris using a BD FACSAria Fusion™ with high-speed sort settings (70 µm nozzle and 70 PSI sheath pressure) and 0.9% NaCl as sheath fluid. The nuclei were identified by PI fluorescence, light scattering, and autofluorescence characteristics (Supplementary Figure 4). A total of 12,600 nuclei were counted and collected in a solution of 4.2 μL phosphate-buffered saline with 0.1% bovine serum albumin.

According to manufacturer’s instructions, the nuclei were loaded into a 10x™ Chromium controller in two batches with 6,300 nuclei each, i.e., two 10x sc-CNV libraries were prepared. In each library, DNA fragments from the same nucleus were ligated with a unique 16-bp barcode sequence (of A/C/G/T). Both libraries were sequenced using Illumina HiSeq3000 in the 2×151 bp paired-end mode, totaling 95 and 124 million read pairs, respectively (61.7 Gb).

### Hi-C library preparation and sequencing

Approximately 0.5 grams of flash-frozen leaf samples of ‘Rojo Pasion’, which were collected from the field, were thawed and fixed with 1% formaldehyde for 30 min at room temperature under vacuum. Subsequently, the in situ Hi-C library preparation was performed according to a protocol established for rice seedlings^44^. The libraries were sequenced on an Illumina HiSeq3000 instrument; in total, around *191*.*2* million pair-end reads were obtained.

### RNA extraction and sequencing

Fruits tissue was collected in the same way for the PacBio sequencing library. Tissue from reproductive buds, vegetative buds, flowers, leaves, bark tissues were collected from the same shoots used for pollen nuclei isolation. RNA was extracted from these tissues using the NucleoSpin® RNA Plant of Macherey-Nagel™ to prepare Illumina libraries.

All libraries were sequenced with Illumina HiSeq 3000 at Max Planck Genome-centre Cologne (MP-GC) in the 150 bp single-end mode, which respectively led to 32.8 (reproductive buds), 28.9 (vegetative buds), 30.2 (flowers), 23.8 (leaves), 18.6 (fruit) and 26.1 (bark) million reads, totaling 24.1 Gb.

### Genome size estimation

After trimming off 10x Genomics barcodes and hexamers from the 61.7 Gb reads of the two 10x sc-CNV libraries, *k*-mer counting (*k*=21) was performed with *Jellyfish*^45^. The *k*-mer histogram was provided to *findGSE*^30^ to estimate the size of the ‘Rojo Pasion’ genome under the heterozygous mode (with ‘*exp_hom=*200’; Supplementary Figure 3).

### Preliminary diploid-genome assembly and curation

With the 19.9 Gb raw PacBio reads of ‘Rojo Pasion’ (Supplementary Figure 2), a preliminary diploid assembly was constructed using *canu*^29^ (with options ‘genomeSize=242500000 corMhapSensitivity=high corMinCoverage=0 corOutCoverage=100 correctedErrorRate=0.105’).

All raw Illumina reads from the 10x libraries were firstly aligned to the initial assembly using *bowtie2*^46^. Then the *purge haplotigs* pipeline was then used to remove haplotigs (i.e., haplotype-specific contigs inflating the true haploid genome) based on statistical analysis of sequencing depth, and identify primary contigs to build up a curated haploid assembly^47^. To reduce the false positive rate in defining haplotigs, each haplotig was blasted to the curated assembly; if over 50% of the haplotig could not be covered by any primary contigs, it was re-collected as a primary contig.

### SNP marker selection

After trimming off 10x barcodes and hexamers, all pooled Illumina reads from the 10x sc-CNV libraries (61.7 Gb) were re-aligned to the curated haploid assembly using *bowtie*2^46^. With 87.2% reads aligned, 989,132 raw SNPs were called with *samtools and bcftools*^48^. Three criteria were used to select potential allelic SNPs (578,209), including i) the alternative allele frequency must be between 0.38 to 0.62; ii) the alternative allele must be carried by 60-140 reads; iii) the total sequencing depth at a SNP must be between 120-280x (as compared with genome-wide mode depth of 208x; Fig. 2b).

### Deletion marker selection and genotyping

The assemblies included 10,452 regions of over 2 kb without SNP marker (total: 110.9 Mb). If the average sequencing depth of such regions was less than or equal to 146x (i.e., the value at the valley between middle and right-most peaks in the sequencing depth distribution; Fig. 2b), it was selected as a deletion-like marker. This revealed 3,253 deletion markers (36.4 Mb), including 237 on contigs without a single SNP marker. The remaining 7,199 regions (74.5 Mb) were defined as conserved (homozygous regions) between two haplotypes (Fig. 2b). For each deletion marker and gamete genome, we assessed the normalized read (*RPKM* value) could of the reads aligned within the deletion using *bedtools*^49^. The genotype at such a deletion marker was initialized as *a* or *n*, where *a* refers to the presence of reads (and therefore relates to the haplotype without the deletion) and *n* refers to the absence of reads (either the deletion haplotype or not having enough information).

### Haplotype phasing and CO identification

Barcodes in the raw reads were corrected using *cellranger*, with which 182.1 million read pairs (51.0 Gb) were clustered into 691 read sets. Reads of each read set were aligned to the curated assembly using *bowtie2*^46^, bases were called using *bcftools*^50^, and a simple bi-marker majority voting strategy was applied to phase the SNPs along each contig (Fig. 2c). After phasing, we identified COs as consistent switches between the haplotypes.

### Ploidy evaluation of single-cell sequencing

For each nucleus, with short read alignment and base calling to the curated assembly, we counted the number of inter-genotype transitions (genotype *a* to *b* and *b* to *a*) at phased SNP markers over all contigs. Correlating this to the number of covered markers revealed two clusters of nuclei (Supplementary Figure 6c). One cluster with 217 nuclei showed that inter-genotype transitions increased linearly with the number of covered markers (while there were high ratios of more than 5 transitions in every 100 markers), which indicated the sequencing data were mixed from more than one nuclei. The other cluster of 445 nuclei (31.2 Gb with 111.4 million read pairs) showed a limited increase (probably due to sequencing errors or markers from repetitive regions), which supported the expected haploid status.

### Imputation of virtual markers at ends of contigs

Let *a* and *b* denote the parental genotypes. The genotype of a nucleus at both ends of a contig (referred to as virtual markers) can be represented by *aa, bb* or *ab* (or *ba*) where *aa*/*bb* indicates an identical genotype along the contig while *ab* (or *ba*) indicates a CO event in the regions of contig. Then we can build up genotype sequences at the two ends of all contigs (with SNP markers) by imputing at all nuclei. For example, given a contig, sequences of *aaaaaa****b****abbbbbbb* (marker 1) and *aaaaaa****a****abbbbbbb* (marker 2) means there is a CO (in bold) at the 7^th^ (of 15) nuclei (Fig. 2c).

### Linkage grouping and genetic mapping

All virtual markers (defined using SNP markers along contigs) were classified into 8 linkage groups (653 contigs: 212.9 Mb) after pairwise comparison of their genotype sequences using *JoinMap4*.*0*^33^ (with haploid population type: HAP; and logarithm of the odds (LOD) values larger than 3.0).

After filtering out contigs with >10% missing nuclei information or nuclei with >10% missing contigs, a high-quality genetic map consisting of 216 contigs (147.3 Mb, corresponding to 622.0 cM; Fig. 3a) was first obtained using regression mapping in *JoinMap* 4.0® with the following settings: LOD larger than 3.0, a “*goodness-of-fit jump”* threshold of 5.0 for removal of loci and a “two rounds” mapping strategy^33^. Genotype sequences imputed at contig ends or deletions (i.e., respective virtual markers) were used to integrate the remaining 723 contigs into the genetic map. For example, given a deletion marker (e.g., *p* and *q* in Fig. 2c-e), if the respective contig had already existed in the genetic map, phasing was only performed at the deletion (according to surrounding phased SNPs); otherwise, phasing plus positioning to the genetic map would be applied. Both operations were based on finding the minimum divergence of the genotype sequence of the marker to that of the other contigs (in the corresponding genetic map). The final genetic map was completed as 891 contigs of 228.0 Mb.

### Haplotype-specific PacBio read separation

PacBio reads (19.9 Gb) were classified based on three major cases after being aligned to the curated assembly using *minimap2*^51^. First, a read covering phased SNP markers was directly clustered into the haplotype supported by the respective alleles in the read. Second, a read covering no SNP markers but overlapping a deletion marker was clustered into the respective genotype based on its phasing with neighboring imputed markers in genetic map. Third, a read in a conserved region was assigned to one of the haplotypes randomly.

### Haplotype assembly and chromosome-level scaffolding

Independent assemblies were performed with sixteen sets of reads, i.e., for every two haplotypes in each of the eight linkage groups using *Flye*^34^ with the default settings.

Using the 891 contigs of the curated assembled that were assigned to chromosomal positions with the genetic mapping, we created a pseudo reference genome, with which the newly assembled contigs were scaffolded using *RAGOO*^52^, leading to chromosome-level assemblies (i.e., those labeled with ‘scaf’ in Fig. 3b).

### Haplotype evaluation

The genotypes of the sixteen assemblies were firstly identified by comparing *k*-mers in each assembly with Illumina WGS of the parental cultivar (*k*=21; Fig. 3c). Although evaluation can always be performed in each linkage group, we combined the eight linkage-group-wise assemblies for ‘Currot’-genotype and the other eight for ‘Orange Red’-genotype, respectively.

After polishing the assemblies respectively with the ‘Currot’-genotype and ‘Orange Red’-genotype PacBio reads using *apollo*^53^, we built up two sets of haplotype-specific *k*-mers from the assemblies, *r*_*C*_ and *r*_*O*_. Correspondingly, a set of ‘Currot’-specific *k*-mers (with coverage from 10 to 60x), *p*_*C*_, was selected from the parental Illumina WGS that did not exist in ‘Orange Red’ short reads (coverage over 1x) but in ‘Rojo Pasion’ pollen short reads (coverage from 10 to 300x); similarly, a set of ‘Orange Red’-specific *k*-mers, *p*_*O*_, was also collected. Then we intersected *r*_*C*_ and *r*_*O*_ with *p*_*C*_ and *p*_*O*_ respectively, leading to four subsets *r*_*C*_⋂ *p*_*C*_, *r*_*C*_⋂ *p*_*O*_, *r*_*O*_⋂ *p*_*C*_, and *r*_*O*_⋂ *p*_*O*_, which were used to calculate average haplotyping accuracy. All *k*-mer processing (counting, intersecting and difference finding) were performed with *KMC*^54^. After haplotype validation, the assemblies were further polished with the respective parental short read alignment using *pilon*^55^ (with options ‘--fix bases --mindepth 0.85’) generating v1.0 of the assemblies. Manual correction of the v.1.0 assemblies was performed according to focal and parental reads to generate assembly v1.1. Finally, *k*-mer-based assembly validation was performed with KAT^35^ and Merqury^36^.

### Genome annotation

We annotated protein-coding genes for each haplotype assembly (v1.0) by integrating evidences from *ab initio* gene predictions (using three tools *Augustus*^56^, *GlimmerHMM*^57^ and *SNAP*^58^), RNA-seq read assembled transcripts and homologous protein sequence alignments. We aligned protein sequences from the database UniProt/Swiss-Prot, *Arabidopsis thaliana* and *Prunus persica* to each haplotype assembly using the tool *Exonerate*^59^ with the options “--percent 60 --minintron 10 --maxintron 60000”. We mapped RNA-seq reads from reproductive buds, vegetative buds, flowers, leaves, fruits (except seeds) and bark tissues, as well as a published Apricot RNA-seq dataset^37^, using HISAT^60^, and we assembled the transcripts using *StringTie*^61^. Finally, we used the tool *EvidenceModeler*^62^ to integrate the above evidence in order to generate consensus gene models for each haplotype assembly.

We annotated the transposon elements (TE) using the tools *RepeatModeler* and *RepeatMasker* (http://www.repeatmasker.org). We filtered the TE related genes based on their coordinates overlapping with TEs (overlapping percent > 30%), sequence alignment with TE-related protein sequences and *A. thaliana* TE related gene sequences (both requiring *blastn* alignment identity and coverage both larger than 30%).

We improved the resulting gene models using in-house scripts. Firstly, we ran a primary gene family clustering using *orthoFinder*^63^ based on the resulting gene models from each haplotype to find haplotype-specific genes. We then aligned these specific gene sequences to the other haplotype using *blastn*^64^ to check whether it was specific because the ortholog was unannotated in the other haplotype. For these potentially unannotated genes (blastn identity > 60% and blastn coverage > 60%), we checked the gene models from *ab initio* prediction around the aligned regions to add the unannotated gene if both the gene model and the aligned region had an overlapping rate larger than 80%. We also directly generated new gene models based on the *Scipio*^65^ alignment after confirming the existence of start codon, stop codon and splicing site. Finally, the completeness of assembly and annotation were evaluated by the *BUSCO*^42^ v4 tool based on 2,326 eudicots single-copy orthologs from OrthoDB v10^66^. A similar process was used to filter for haplotype-specific genes (Supplementary Tables 2-3). Finally, a genome annotation lift-over was performed from v1.0 to v1.1 using liftoff^67^ with default parameters.

### Genome assembly comparison

All genome assemblies, including ‘Rojo Pasion’ haplotypes, ‘Chuanzhihong’ apricot (*Prunus armeniaca*)^37^, Japanese apricot (*Prunus mume*)^38^ and ‘Lovell’ peach (*Prunus persica*)^39^, were aligned to each other using *nucmer* from the *MUMmer4*^68^ toolbox with parameters ‘-max -l 40 -g 90 -b 100 -c 200’. The alignments were further filtered for alignment length (>100 bp) and identity (>90%), with which structural rearrangements and local variations were identified using *SyRI*^23^. To follow the nomenclature of the Prunus community, the ‘Rojo Pasion’ chromosomes were numbered according to the numbering in ‘Lovell’ peach^39^.

### Hi-C data analysis

We used *ALLHiC*^8^ and *SALSA2*^40^ for phasing and scaffolding. All 191.2 million Hi-C read pairs were aligned (using *BWA* version 0.7.15-r1140) to the haplotype-resolved unitigs assembled by *Canu*. Only uniquely mapped read pairs were selected using *filterBAM_forHiC*.*pl* from the *ALLHiC* package. The selected alignments were used as input for *ALLHiC_partition* (“ALLHiC_partition -b clean.bam -r unitigs.fa -e GATC -k 19”) and *SALSA2* (“python run_pipeline.py -a unitigs.fa -l unitigs.fa.fai -g unitigs.gfa -m yes -b alignment.bed -e GATC -o SALSA2_out -i 8”, where the file alignment.bed was generated and sorted from clean.bam using *bedtools bamtobed* (version v2.29.0) and unitigs.gfa was collected from the *Canu* output). For *ALLHiC*, we had to set group number as 19 to get 16 linkage groups (of chromosome-level size), and 3 smaller groups below 2.5 Mb, which were not considered further. We continued with *ALLHiC* pipeline as it provided more accurate than those from *SALSA2*. The subsequent pipeline of *ALLHiC* were run by default except for using “-RE GATC” in the “allhic extract” command. For comparison, we also aligned all raw Hi-C reads to haploid assemblies generated by gamete binning, and selected the uniquely mapped read pairs as described above. Hi-C maps were visualized using *ALLHiC_plot* at 300-500 kb resolution. Alignments of *ALLHiC* and gamete binning based assemblies were obtained using *minimap2* and dot plot was drawn with script *pafCoordsDotPlotly*.*R* at *https://github.com/tpoorten/dotPlotly*.

### Crossover identification

All 220 million pollen nuclei-derived short read pairs were pooled and aligned to the ‘Currot’-genotype assembly, from which 739,342 SNP markers were defined with an alternative allele frequency distribution of 0.38 to 0.62 and alternative allele coverage of 50 to 150x. Then, short reads of 445 nuclei were independently aligned to the ‘Currot’-genotype assembly using *bowtie2*^46^ and bases were called using *bcftools*^50^. Finally, *TIGER*^69^ was used to identify COs. The landscape of COs from 369 nuclei with a sequencing depth over 0.1x was calculated within 500 kb sliding windows along each chromosome at a step of 50 kb (Fig. 5), where for each window, the recombination frequency (*cM/Mb*) was defined as *C*/*n*/(*w*/10^6)* 100%, where *C* is the number of recombinant nuclei in that window, *n* is the total number of nuclei (369) and *w* is the window size. *SNP/Mb* and *gene/Mb* were calculated for the same windows as *x/*(*w*/10^6), where *x* was the count of the feature in the respective window.

## Acknowledgements

The authors would like to thank Antonio Molina and José Egea for kindly providing plant material, Saurabh Pophaly for help in transferring the read data to public servers, Detlef Weigel for supportive guidance, and Kristin Krause and Vidya Oruganti for useful discussions and comments for improving the manuscript. This work was funded by the “Humboldt Research Fellowship for Experienced Researchers” (Alexander von Humboldt Foundation) (J.A.C.), the Marie Skłodowska-Curie Individual Fellowship PrunMut (789673) (J.A.C.), the Deutsche Forschungsgemeinschaft (DFG, German Research Foundation) under Germany’s Excellence Strategy – EXC 2048/1– 390686111 (K.S.), and the European Research Council (ERC) Grant “INTERACT” (802629) (K.S.). C.K. acknowledges the ISAC SRL Emerging Leaders Program.

## Author contributions

J.A.C, H.S. and K.S. designed the project. J.A.C., B.H., K. F.-D., C.K., D.R., M.R., N.W. and C.L. performed wet-lab experiments. H.S., J.A.C, M.G., and W-B.J. performed all data analysis. J.A.C., H.S. and K.S. wrote the manuscript with input from all authors. All authors read and approved the final manuscript.

## Competing interests

The authors declare no competing interests.

## Data availability

Data supporting the findings of this work are available within the paper and its Supplementary Information files. Read data sequenced from two 10x sc-CNV libraries, two Hi-C libraries, one PacBio library from ‘Rojo Pasion’, two Illumina libraries for ‘Currot’ and ‘Orange Red’ that support the work in this study as well as the haploid assemblies and annotations generated are available in European Nucleotide Archive (ENA) under accession number “PRJEB37669”. Data was uploaded to ENA using EMBLmyGFF^70^. All other relevant data are available upon request.

## Code availability

Customs scripts supporting this work are available at https://github.com/schneeberger-lab/GameteBinning.

## Notes

### Competing Interest Statement

The authors have declared no competing interest.

### Summary of Updates

Comparison of gamete binning to Hi-C added; k-mer analysis to validate the assemblies added; author list and affiliations updated; Fig. 3e and Fig. 4. added; Supplementary Information updated.

