## Supplementary_Information for "Chromosome-level and haplotype-resolved genome assembly enabled by high-throughput single-cell sequencing of gamete genomes"

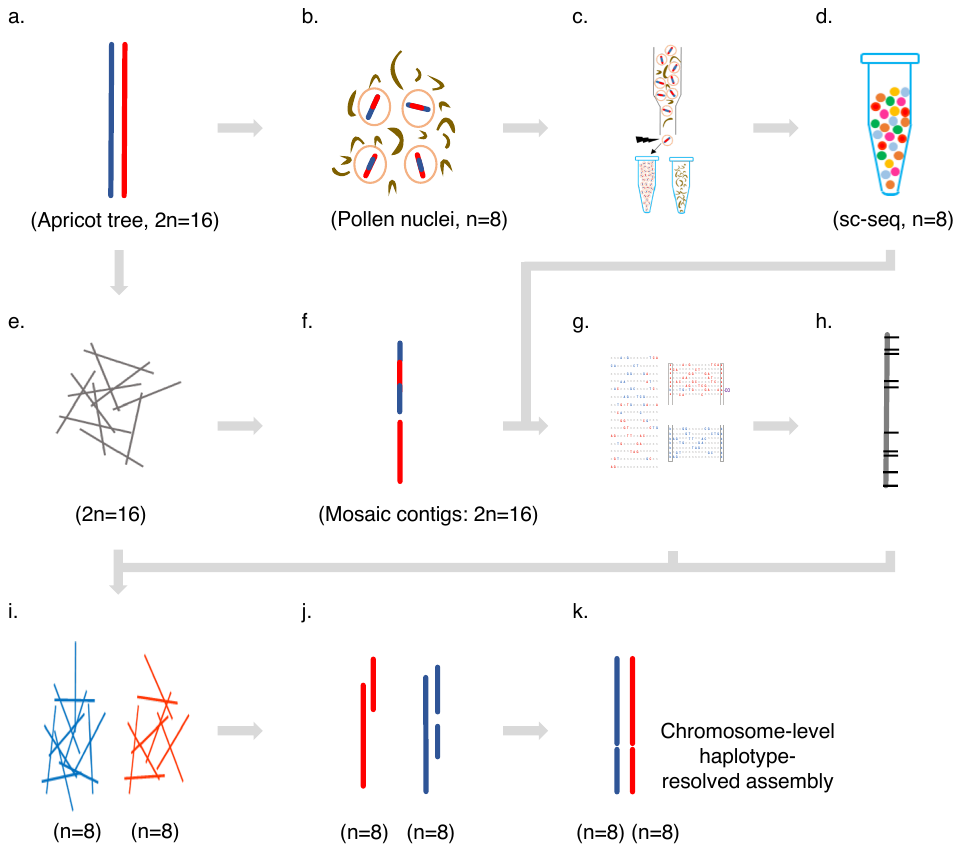

**Supplementary Figure 1. Flowchart of gamete binning in haploid assembly of apricot (*Prunus armeniaca* cultivar ‘Rojo Pasion’).** a. Heterozygous diploid apricot tree genome (represented by one chromosome pair; *n* gives the ploidy level). b. Isolation of ~500,000 pollen nuclei (mixed with debris) from ten flowers. c. Nuclei selection with flourescence activated cell sorting. d. Single-pollen nuclei genome sequencing (sc-seq: 10x Chromium System^TM^ plus Illumina Hi-seq 3000). e. Long read sequencing of somatic tissue (PacBio). f. Initial mosaic haploid assembly using the full set of long reads. g. Variant phasing using sc-seq and the mosaic haploid assembly. h. Contig grouping and linkage mapping. i. Separation of long reads to haplotypes in linkage groups (8x2 sets) using phased variants. j. Independant assembly of each haplotype. k. Scaffolding contigs (from j) using the gamete genome-derived genetic map (in h), leading to chromosome-scale haplotype-resolved assembly.

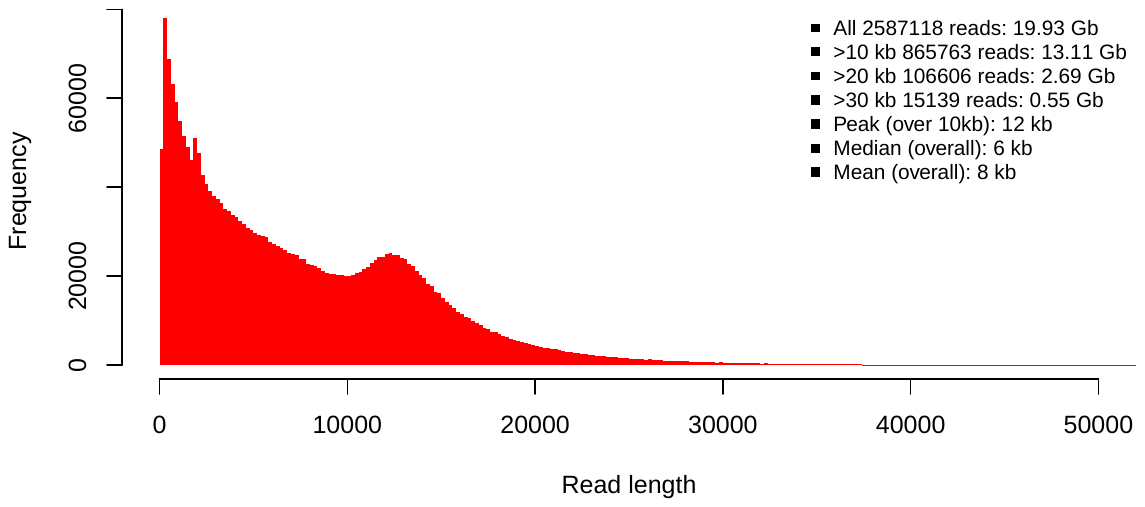

**Supplementary Figure 2. Size distribution of long reads (PacBio).** As given, there were 2,587,118 reads totaling 19.93 Gb.

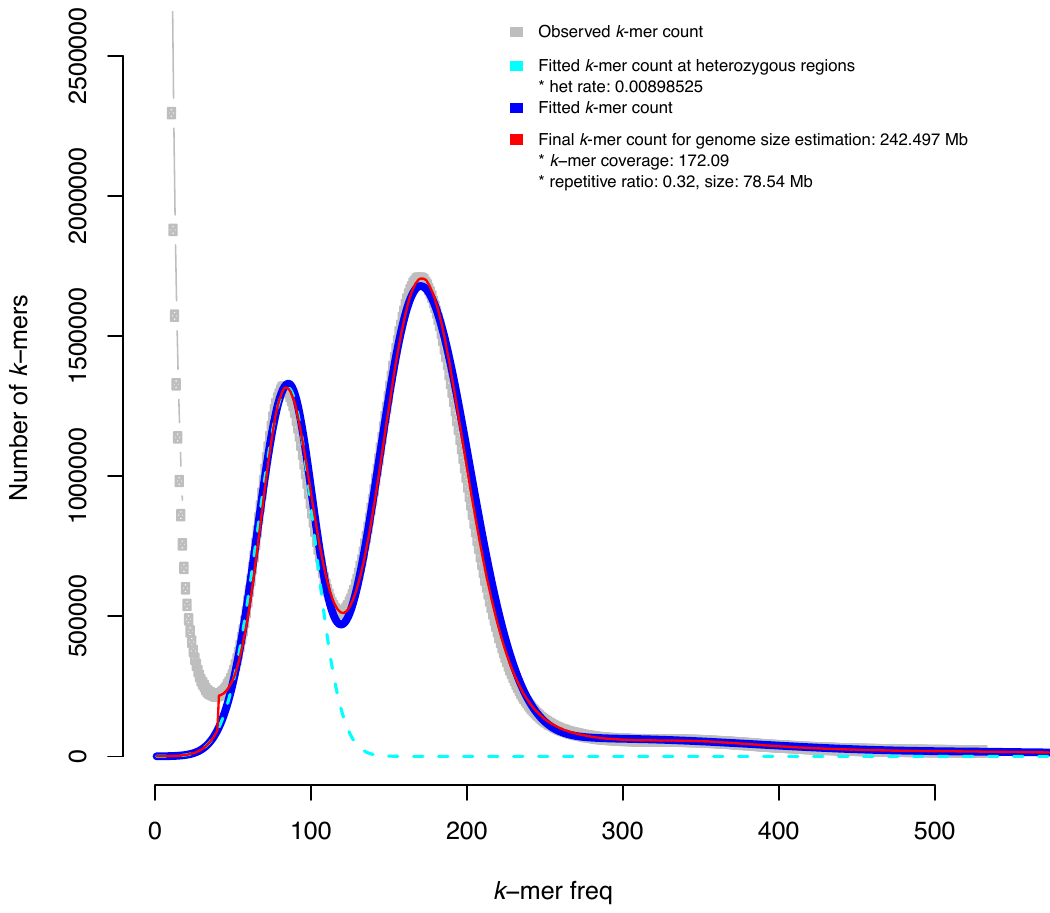

**Supplementary Figure 3. Genome size estimation with *k*-mers of pooled Illumina reads from pollen nuclei (*k*=21).** Genome size was determined as 242.5 Mb and heterozygosity was 0.9%. Note: before *k*-mer counting, the 16 bp 10x barcode and a haxmer in the beginning of each read 1 were trimmed off.

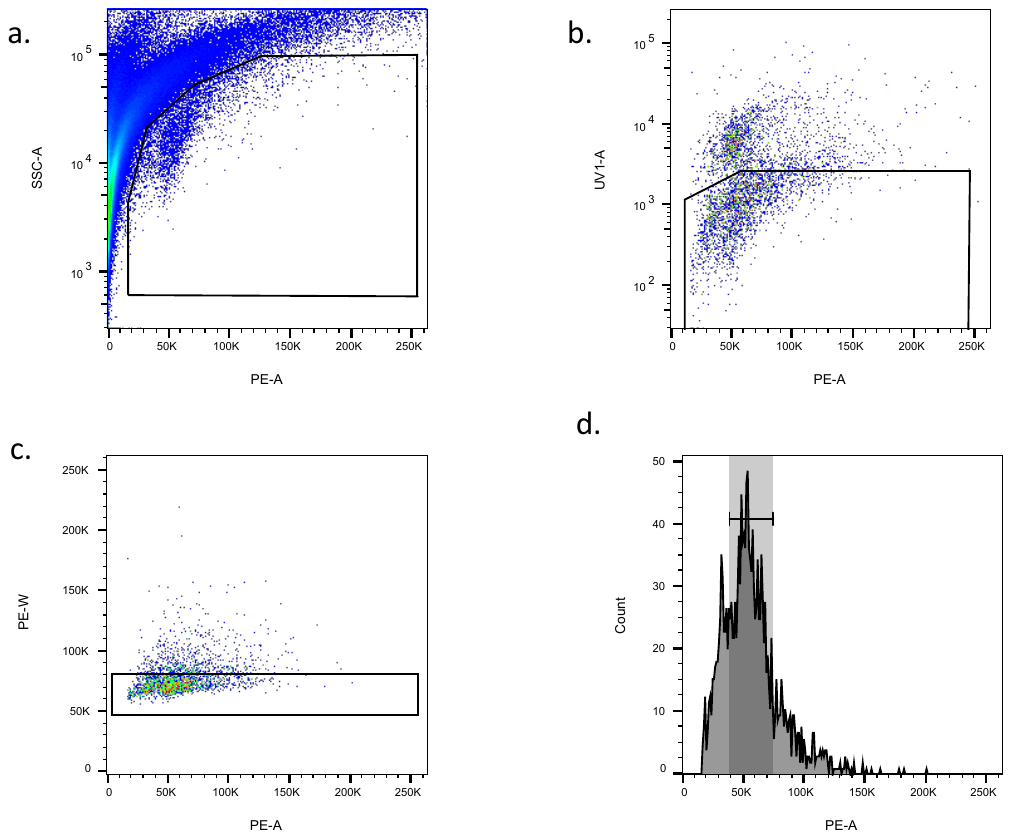

**Supplementary Figure 4. Illustration of flow cytometry sorting of pollen nuclei.** Individual pollen nuclei were subsequentially identified by and sorted on the following characteristics: **a**. Light scatter (SSC-A; log scale) and propidium iodide (PI) fluorescence (PE-A; linear scale; 561 nm excitation; 586/15 nm emission). **b**. Low UV/violet autofluorescence (UV1-A; log scale; 355 nm excitation; 450/50 nm emission) and PI fluorescence. **c**. Singlets (pulse width, PE-W; linear scale) and propidium iodide fluorescence. Colors (a, b, c) reflect levels of nuclei density (blue < green < yellow < orange < red) according to “pseudo-color density plots” provided by FlowJo vX^TM^ software. **d**. High but modal PI fluorescence (PE-A). Finally, shaded rectangular indicates selected nuclei.

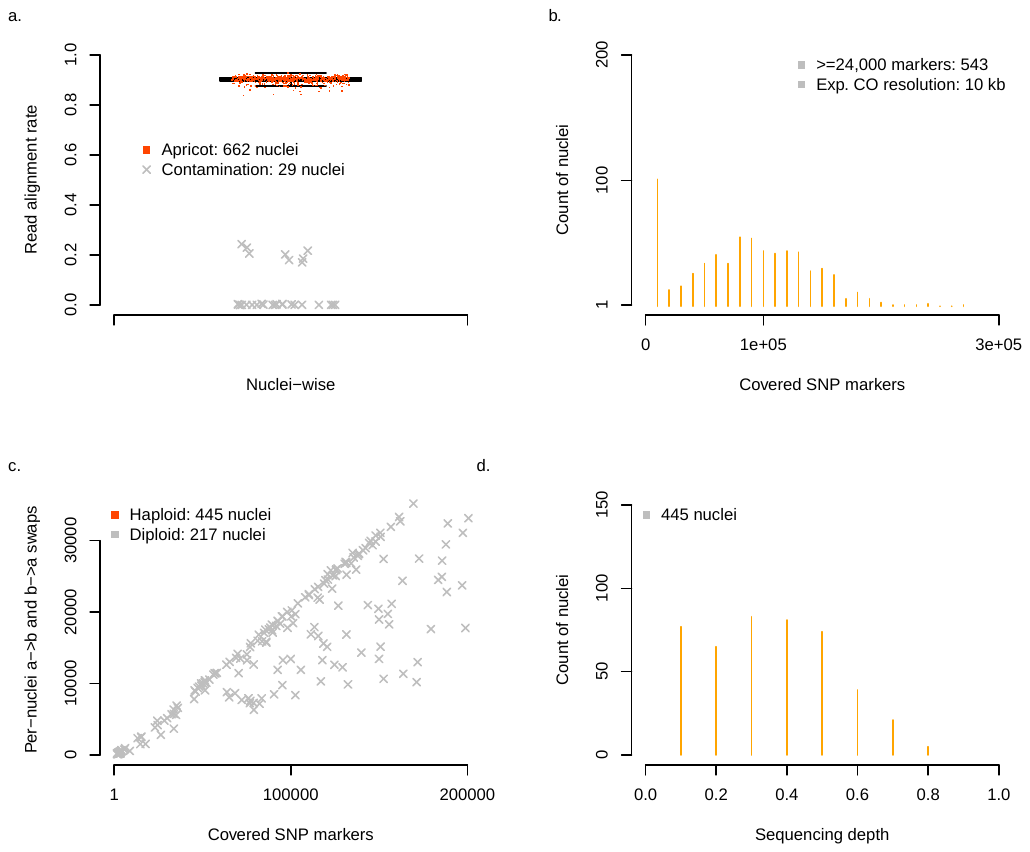

**Supplementary Figure 5. Characterization and selection of pollen nuclei sequencings. a**. Read alignment rate identified 29 nuclei contaminations (such as from thrips genomes) which were removed from further analysis. The average alignment rate of 662 nuclei from apricot was 90%. **b**. Coverage of SNP markers in single nuclei. There were 543 of 662 nuclei with over 24,000 SNP markers covered, indicating an expected crossover (CO) resolution of 10 kb. **c.** Identification of haploid pollen nuclei. Let *a* and *b* denote two genotypes. X-axis: total number of SNP markers covered in a sample. Y-axis: total number of transitions from *a* to *b* and *b* to *a* in a sample. The association revealed that 445 samples were haploid and thus selected for further analysis, while 217 samples were with reads from more than 1 nuclei (and thus removed; main text: Methods). **d**. Sequence depth of 445 selected haploid nuclei, among which 369 nuclei were above 0.1x, and later used for CO landscape analysis.

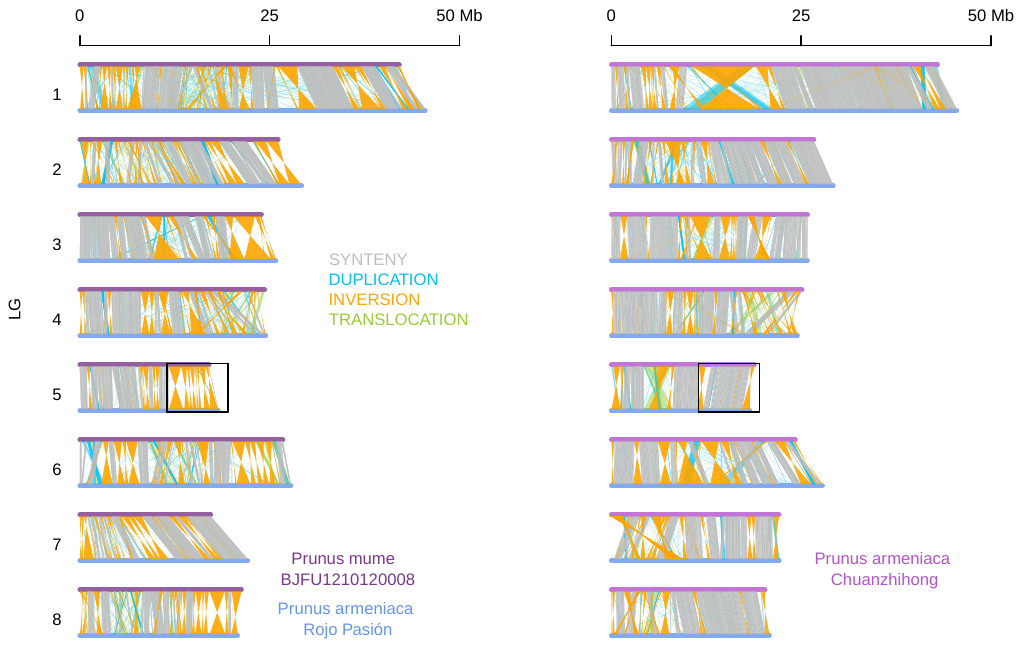

**Supplementary Figure 6. High-scaffolding accuracy reflected by synteny to closely-related species: ‘Chuanzhihong’ apricot (*Prunus armeniaca*)^1^ and Japanese apricot (*Prunus mume*)^2^.** Substantial structural variations (SVs, including duplications, inversions and translocations) were found between genome assemblies of ‘Rojo Pasion’ (using the haplotype inherited from parental ‘Currot’ as the representative) and these species (as well as peach in Fig. 2d in main text). However, the SVs were hardly identified simultaneously, for example, within a region (in black box) on LG 5, while there were many inversions found between ‘Rojo Pasion’ and Japanese apricot, most of them were not found in the comparison between ‘Rojo Pasion’ and ‘Chuanzhihong’. These comparisons indicated either the differences from the respective species were real or that the assembly generated in this work was of high quality.

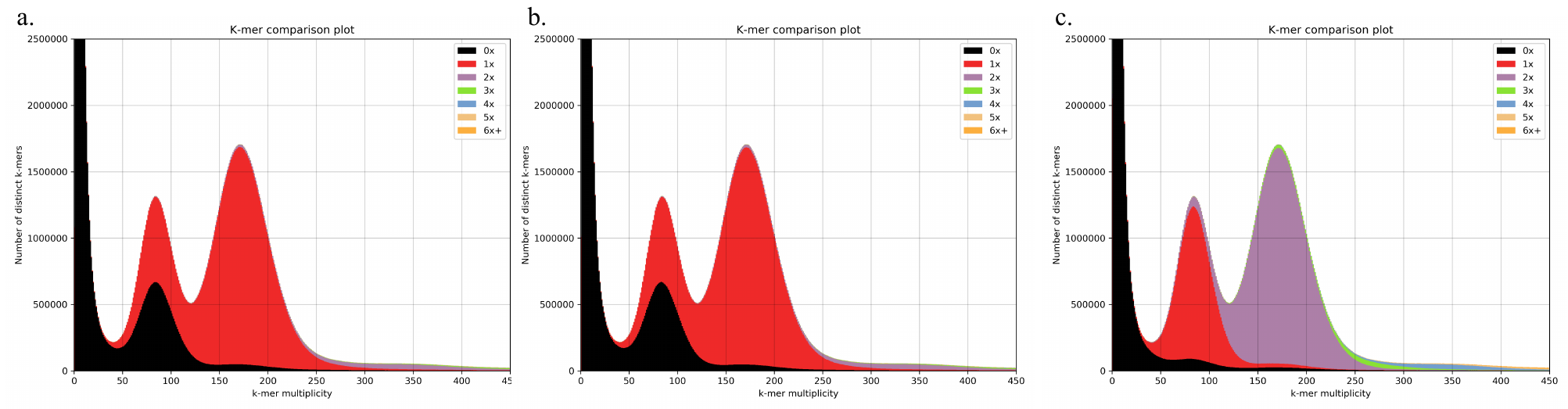

**Supplementary Figure 7. *k*-mer comparison plot for genome assembly evaluation by KAT3**^3^**.** Parental (‘Rojo Pasion’) read *k*-mer frequency versus Orange Red haplotype (*a*), Currot haplotype (*b*) and Rojo Pasion (Orange Red and Currot haplotypes merged, *c*) copy number stacked histograms. Read content in black is absent from the assembly, red occurs once, purple twice, etc. *k*-mer spectra show an error distribution under 45x, heterozygous content around 85x and homozygous content around 170x. *a* and *b*, show correct single haplotype mosaics with minimal level of artificial duplication. *c*, shows the separated haplotypes merged producing the duplication of content in the homozygous regions, allowing the full representation of heterozygous content. No significant artificial duplication was found (purple signal in 85x heterozygous peak and green signal in 170x homozygous peak).

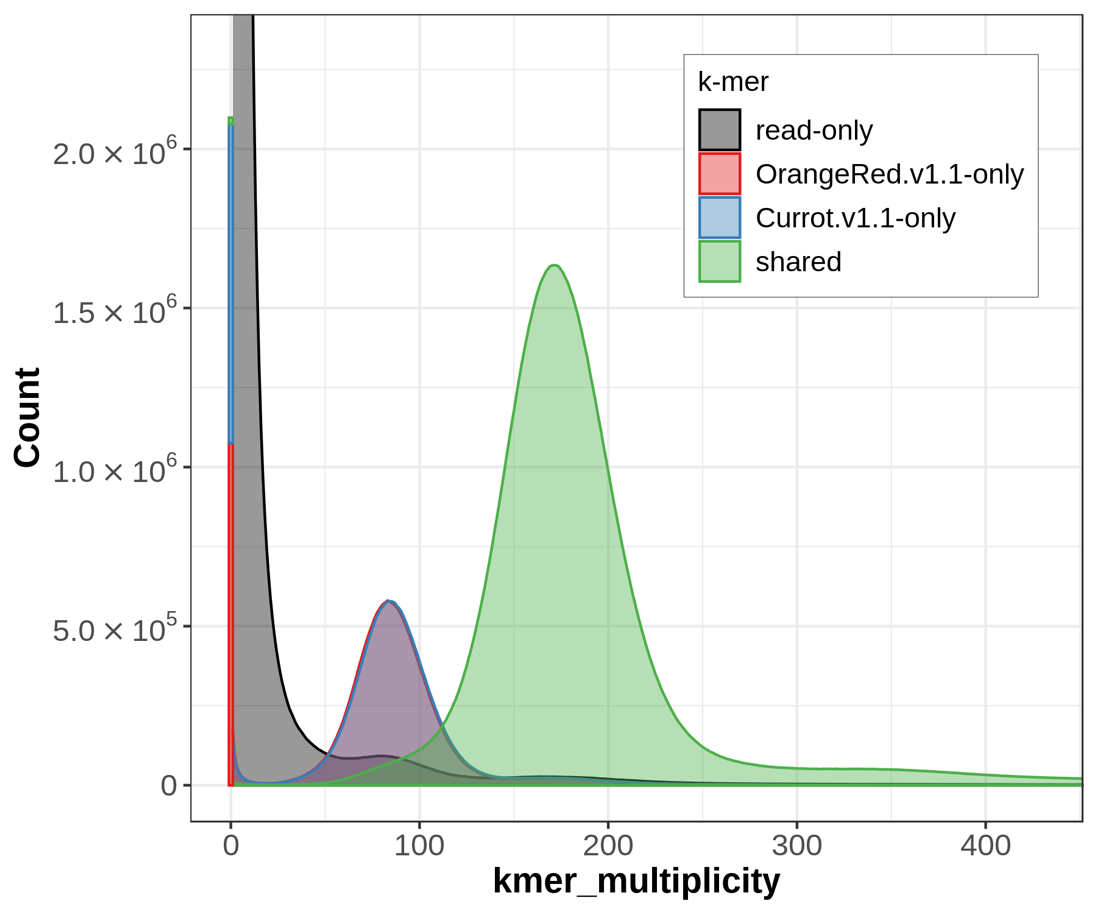

**Supplementary Figure 8. Assembly spectrum plots for evaluating *k*-mer completeness by Merqury^4^.** *k*-mers are colored by their presence in the ‘Rojo Pasion’ reads and OrangeRed/Currot/RojoPasion (shared) assemblies. The small signal (~7%, around 85x in the black curve, representing 1-copy k-mers found only in the reads) corresponds to the fraction of heterozygous variants missing from the assembly (93.4% of Pac-Bio reads were separated) plus erroneous *k*-mers.

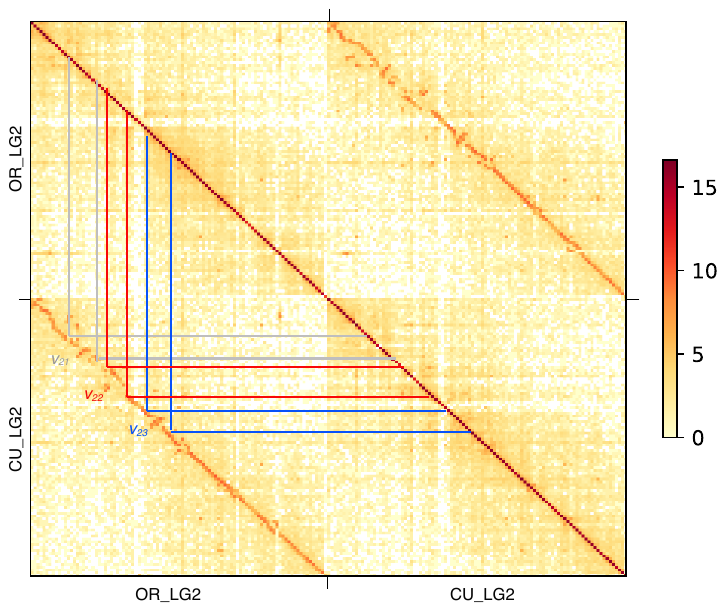

**Supplementary Figure 9.** Hi-C contact along gamete binning based assemblies for chromosome 2 related to both haplotypes of Currot (CU) and Orange Red (OR) (with bin size or resolution of 300 kb). Variants spanning over 500 kb are labelled as v_xy_, where *x* denotes the chromosome number and *y* the number of the large variant in the chromosome, and validate the structural rearrangements predicted by *SyRI* in the gamete-binning assemblies (Figure 5).

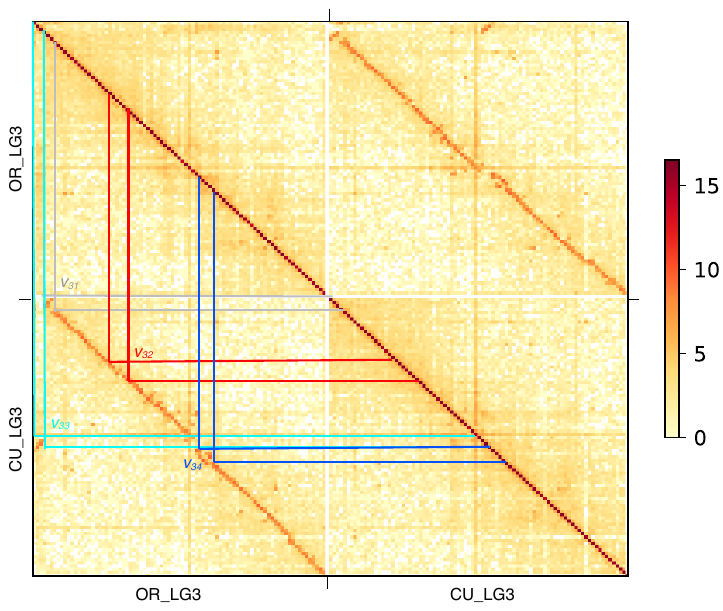

**Supplementary Figure 10.** Hi-C contact along gamete binning based assemblies for chromosome 3 related to both haplotypes of Currot (CU) and Orange Red (OR) (with bin size or resolution of 300 kb). Variants spanning over 500 kb are labelled as v_xy_, where *x* denotes the chromosome number and *y* the number of the large variant in the chromosome, and validate the structural rearrangements predicted by *SyRI* in the gamete-binning assemblies (Figure 5).

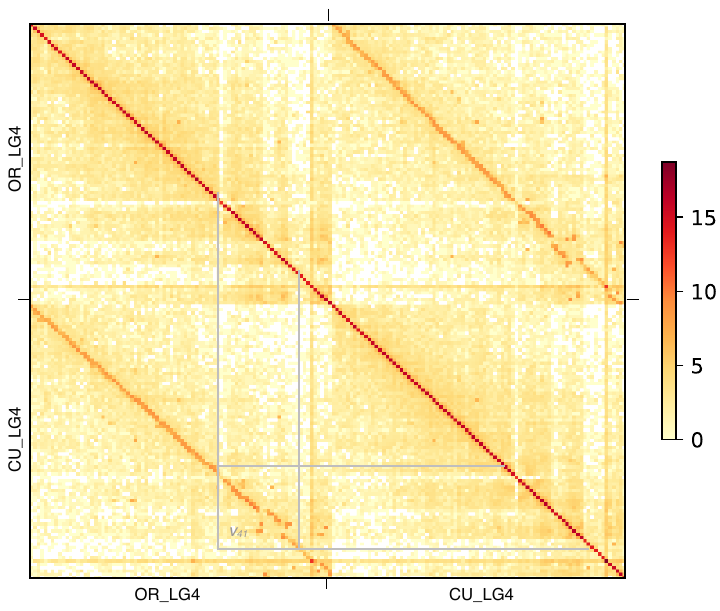

**Supplementary Figure 11.** Hi-C contact along gamete binning based assemblies for chromosome 4 related to both haplotypes of Currot (CU) and Orange Red (OR) (with bin size or resolution of 300 kb). Variants are labelled as v_xy_, where *x* denotes the chromosome number and *y* the number of the large variant in the chromosome, and validate the structural rearrangements predicted by *SyRI* in the gamete-binning assemblies (Figure 5). v_41_ is shown as an example of Hi-C validation of a set of small rearrangements spanning less than 500 kb.

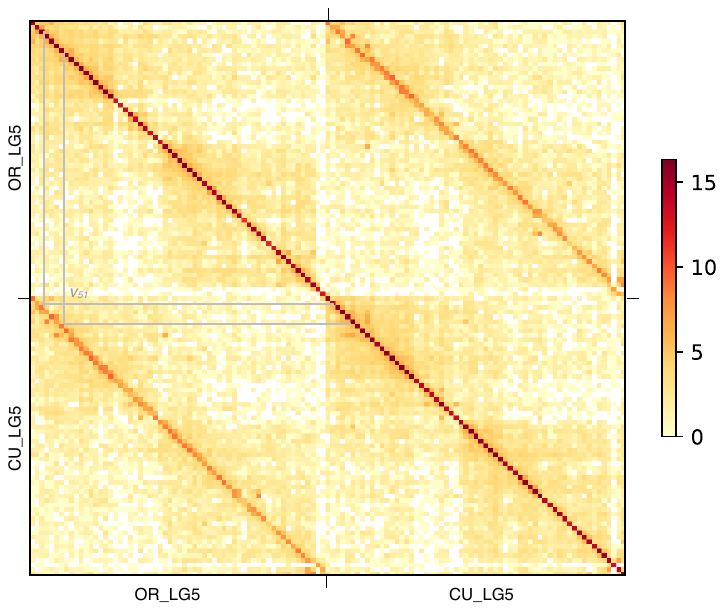

**Supplementary Figure 12.** Hi-C contact along gamete binning based assemblies for chromosome 5 related to both haplotypes of Currot (CU) and Orange Red (OR) (with bin size or resolution of 300 kb). Variants spanning over 500 kb are labelled as v_xy_, where *x* denotes the chromosome number and *y* the number of the large variant in the chromosome, and validate the structural rearrangements predicted by *SyRI* in the gamete-binning assemblies (Figure 5).

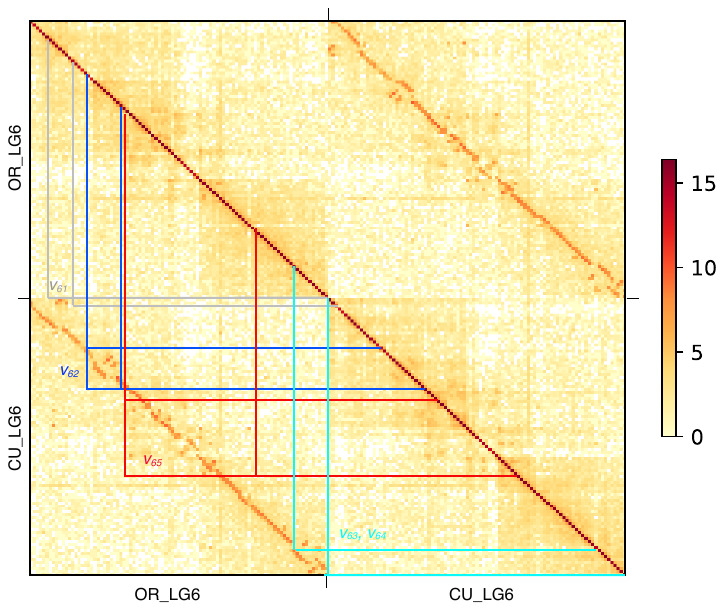

**Supplementary Figure 13.** Hi-C contact along gamete binning based assemblies for chromosome 6 related to both haplotypes of Currot (CU) and Orange Red (OR) (with bin size or resolution of 300 kb). Variants spanning over 500 kb (except v_65_) are labelled as v_xy_, where *x* denotes the chromosome number and *y* the number of the large variant in the chromosome, and validate the structural rearrangements predicted by *SyRI* in the gamete-binning assemblies (Figure 5). v_65_ is shown as an example of Hi-C validation of a set of small rearrangements spanning less than 500 kb.

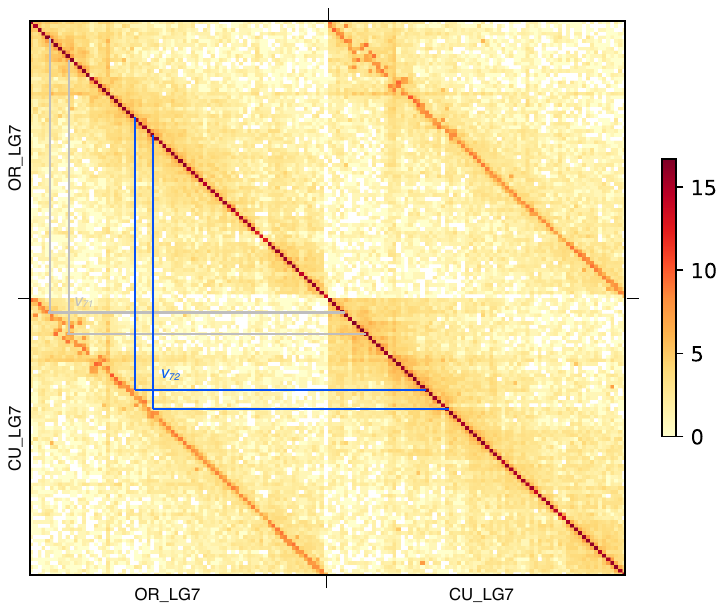

**Supplementary Figure 14.** Hi-C contact along gamete binning based assemblies for chromosome 7 related to both haplotypes of Currot (CU) and Orange Red (OR) (with bin size or resolution of 300 kb). Variants spanning over 500 kb are labelled as v_xy_, where *x* denotes the chromosome number and *y* the number of the large variant in the chromosome, and validate the structural rearrangements predicted by *SyRI* in the gamete-binning assemblies (Figure 5).

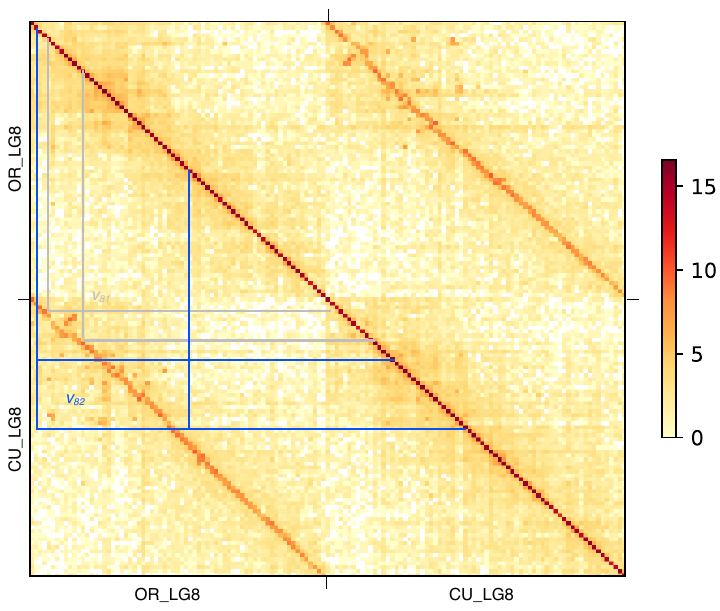

**Supplementary Figure 15.** Hi-C contact along gamete binning based assemblies for chromosome 8 related to both haplotypes of Currot (CU) and Orange Red (OR) (with bin size or resolution of 300 kb). Variants spanning over 500 kb (except v_82_) are labelled as v_xy_, where *x* denotes the chromosome number and *y* the number of the large variant in the chromosome, and validate the structural rearrangements predicted by *SyRI* in the gamete-binning assemblies (Figure 5). v_82_ is shown as an example of Hi-C validation of a set of small rearrangements spanning less than 500 kb.
